# LEMMIv2: Benchmarking Framework for Metagenomic and 16S Amplicon Profilers with a Catalogue of Evaluated Tools

**DOI:** 10.1101/2025.03.06.641904

**Authors:** M. Seppey, A. Benavides, M. R. Berkeley, M. Manni, E. M. Zdobnov

## Abstract

Sequencing has transformed microbial studies, enabling metagenomic analysis of microbial communities without the need for culturing or prior knowledge of sample composition. The essential analysis of the primary sequencing reads, however, is complex and led to the development of diverse computational strategies. The plethora of available methods and their numerous parameters poses a practical challenge for practitioners and creates a visibility barrier for developers of novel approaches. In addition to technical limitations related to user computing environment, algorithmic solutions, and their scalability, there are critical considerations regarding the reference database, i.e. the knowledge against which the data is interpreted, as well as the target of the analysis, expected sample composition, and peculiarities of read data from different sequencing platforms. To facilitate informed decision-making, we introduced the LEMMI platform for continuous benchmarking of software tools for metagenomic analyses, where developers can receive impartial benchmarks for method publication and users benefit from a standardized and benchmarked catalogue of tools. Here we present developments of LEMMI version 2, including assessments of different target scenarios, long- and short-read sequencing data, alternative taxonomies, and a standalone pipeline (https://lemmi.ezlab.org). In addition to LEMMI, which focuses on shotgun metagenomic profiling, we extended this approach to bacteria profiling with 16S amplicon sequencing (https://lemmi16S.ezlab.org).

## Introduction

Next-generation sequencing has become the state-of-the-art method for studying microorganisms, enabling the profiling of entire microbial communities through shotgun metagenomics, including species that have not been cultivated and without prior knowledge of the sample’s contents. Over the past decade, data production has increased exponentially. In response, researchers have developed numerous computational methods to assign taxonomic labels to sequencing reads more efficiently and at scale, while improving classification quality (e.g., Wood et al., (2019), Blanco-Míguez et al., (2023)). Along with the growing volume of data, the evolution of sequencing technologies has driven innovation in the field. Illumina platforms, which produce short reads, remain the most widely used due to their accuracy and throughput. However, novel long-read sequencing technologies from PacBio and Oxford Nanopore (ONT) are increasingly being used for metagenomics (Agustinho et al., 2024), leading to the development of computational methods fine-tuned to their specific characteristics.

Individual research groups and larger communities have surveyed and evaluated these tools to help users make informed decisions. Self-benchmarking has also become a standard practice when publishing new tools. While informative, one-off benchmarking papers provide snapshots that quickly become outdated. When authors introduce new metrics and innovative dataset designs, they often make comparisons with existing data difficult or impossible. Given the proliferation of benchmarking methods in various areas of genomics, including metagenomic profiling, there is a need for more sustainable and reproducible approaches in the long term (Mangul et al., 2019).

In this context, we developed the LEMMI benchmarking platform for metagenomics profilers (Seppey et al., 2020), available at https://lemmi-v1.ezlab.org/. Our objective was to offer a framework for continuous assessment of candidate tools under stable conditions, using public data to simulate microbial communities and Illumina datasets, while controlling the reference material available to the tools. This platform can incorporate tools that did not exist when the benchmark was first designed, maintaining a consistent challenge for newcomers. The benchmark results are presented on an interactive web app. The platform ensures access to the evaluated methods through publicly available software containers, which persist even if developers discontinue the maintenance of the tools. Since launching the platform, we have enabled submissions of new tools via software containers, provided they followed a specific design procedure that allowed our team to conduct the evaluation. While this approach fulfilled our initial goal of maintaining an incremental benchmark, we later aimed to provide a LEMMI framework that allows developers and users to run evaluations independently. This would enable them to replicate our analyses, design their own benchmarks, and more easily prepare their tools to be compatible with the LEMMI procedure we run. This manuscript presents version 2 of LEMMI (LEMMIv2, https://lemmi.ezlab.org/) for metagenomics profilers, highlighting key innovations such as a standalone pipeline and the addition of long-read technology. This version builds on our learnings from the initial release and represents a complete refactoring of both the benchmarking methodology and the results presentation interface.

The second objective of this manuscript is to introduce LEMMI16S (https://lemmi16S.ezlab.org/), a dedicated LEMMI benchmark for amplicon sequencing. Amplicon sequencing remains popular for profiling microbial communities due to its cost-effectiveness, well-established protocols, and reduced computational power requirements compared to shotgun metagenomics. In amplicon sequencing, the entire 16S ribosomal RNA gene or subsets of its hypervariable regions (the 16S rRNA gene contains nine hypervariable regions) are used for microbial identification. Although traditionally considered limited to genus or species-level resolution, new developments are challenging this limitation (Johnson et al., 2019; Buetas et al., 2024). An amplicon sequencing analysis typically includes preliminary steps to remove primers, low-quality reads, chimeras, and other PCR and sequencing artifacts. Subsequently, the remaining reads are either clustered into Operational Taxonomic Units (OTUs), grouping sequences at 95% or 97% nucleotide identity as a proxy for genus or species level (Schloss & Handelsman, 2005), or arranged into Amplicon Sequence Variant (ASV) tables through denoising and correction procedures without clustering (Callahan et al., 2016). Finally, these resulting OTU groups or ASV tables are used in classification methods to assign taxonomic levels. These tasks and their advancements have led to the development of new software suites, which also require thorough evaluation. LEMMI16S addresses this need by providing a standardised benchmark for assessing the performance and accuracy of various amplicon sequencing tools across various scenarios.

## Results

### The LEMMIv2/16S framework

#### A workflow that promotes sharing while centralizing knowledge

The LEMMIv2/16S workflow centres around an online catalogue of evaluated tools that has been significantly refactored compared to LEMMIv1. In the first version, datasets for evaluation had to be manually generated or sourced from public datasets. In contrast, the LEMMIv2/16S framework is designed to generate new benchmarking material on demand, using public data to simulate scenarios that mimic real-life problems. These scenarios, referred to as LEMMI instances, help users identify the most relevant tools for their needs. Users can set up multiple parameters to generate these instances, including the target clade (PROK for prokaryotes, EUK for eukaryotes, and VIR for viruses), the taxonomic composition of the microbial community, the sequencing technology, and the taxonomy (NCBI (Schoch et al., 2020) or the Genome Taxonomy Database (GTDB; Parks et al., 2018), a new feature in LEMMIv2). Based on these definitions, *in-silico* reads are produced. For example, the “2023_12_PROK_NCBI_clean_v220” instance in the LEMMIv2 catalogue includes 2 million reads (Illumina HiSeq 2500, read length 150bp) representing 155 species, simulating a clinical sample from dental plaque. The “Various human pathogens, regions V1-V2” instance in the LEMMI16S catalogue includes 845,000 reads (Illumina MiSeq v3, read length 200bp) from the V1-V2 region of the 16S gene, representing 12 species of various human pathogens. As in LEMMIv1, each scenario corresponds to an online report with metrics such as precision, recall, runtime, and memory consumption of the tools benchmarked. Alongside benchmark results, the catalogue provides links to obtain the tools and the necessary information to understand and use them. A major improvement in LEMMIv2/16S is the introduction of a standalone pipeline, similar to the one used to create the online catalogue, which method developers can use to set up local evaluations. This makes it much easier to create a LEMMI-compliant software container for their tools compared to LEMMIv1. Developers can generate their own LEMMI instances or easily obtain existing datasets or configuration files to replicate an existing benchmark. Any tool container compatible with LEMMI can be obtained through the LEMMI website or a third-party repository and used in personal benchmarks with minimal effort. Finally, developers who choose to publish their method can have their LEMMI-compliant container directly evaluated and included in the ongoing public benchmark maintained by the LEMMIv2/16S development team (Figure 1).

**FIGURE 1.**
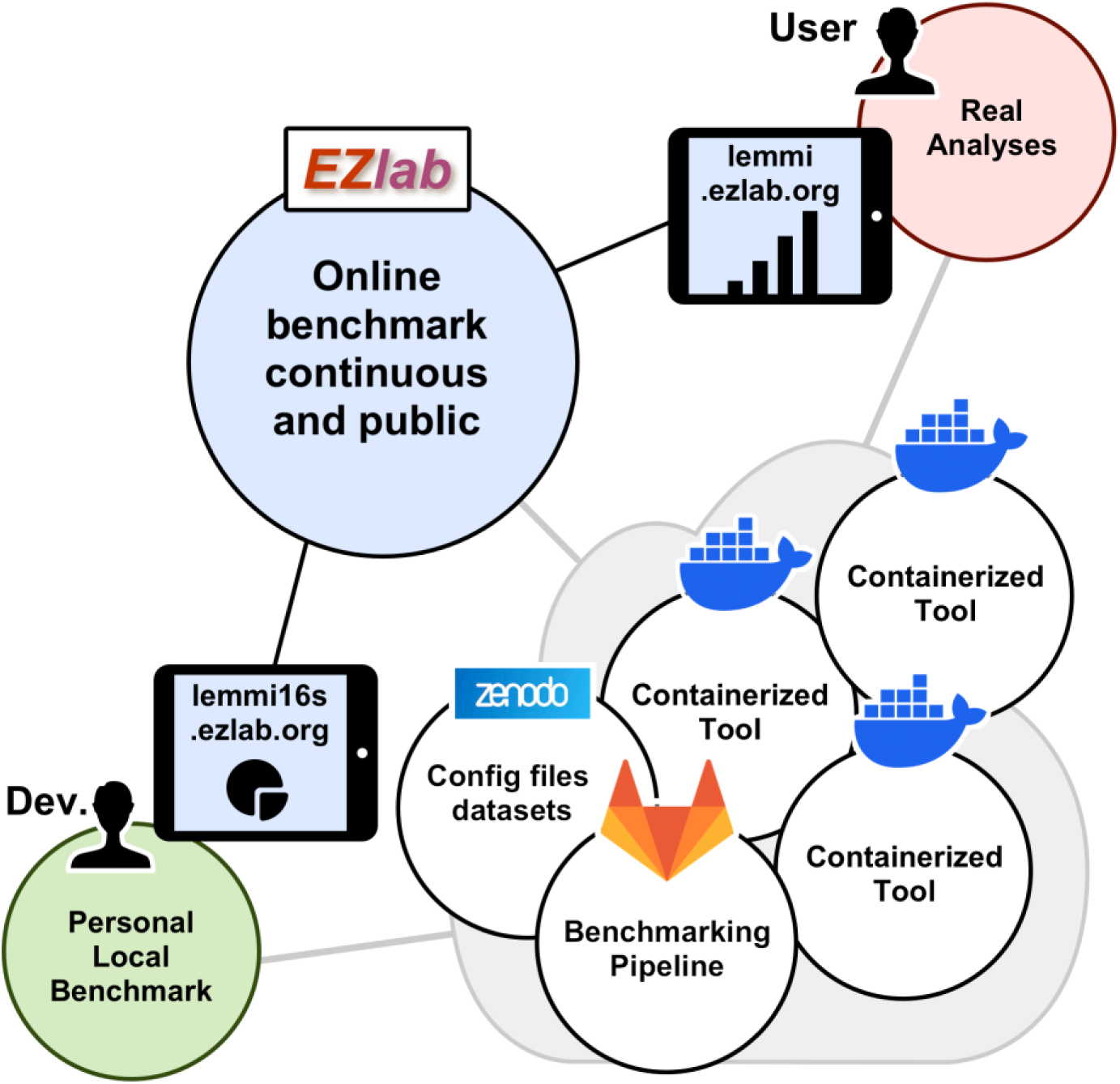
The LEMMIv2/16S workflow. The benchmarking pipeline is publicly available and can be used by any tool developer or user in their own computing environment. It is also used by the LEMMIv2/16S development team to maintain a continuously updated catalogue of evaluated tools that can be consulted online. All methods are packaged in software containers that can be shared in public repositories and are available for re-evaluation or use in real-world analyses. Additional compatible datasets can be shared on Zenodo and reused with the pipeline.

#### An online catalogue of tools evaluated across a wide range of samples

The landing pages of LEMMI16S (Figure 2A) and LEMMIv2 (Figure 2B) showcase a variety of benchmark instances available in the current catalogue. These instances represent a range of scenarios, each differing in terms of community composition, simulated sequencing technologies, and reference taxonomies. In the LEMMIv2 catalogue, users can explore instances that simulate clinical samples, reflecting bacterial communities found in human patients, as well as microbial communities from environmental niches such as alpine lakes. Additionally, users can examine scenarios utilising short reads, as with LEMMIv1, alongside long reads simulated with Oxford Nanopore Technology. These instances are analysed using the NCBI taxonomy, as in the previous version, or the GTDB taxonomy as an alternative.

**FIGURE 2.**
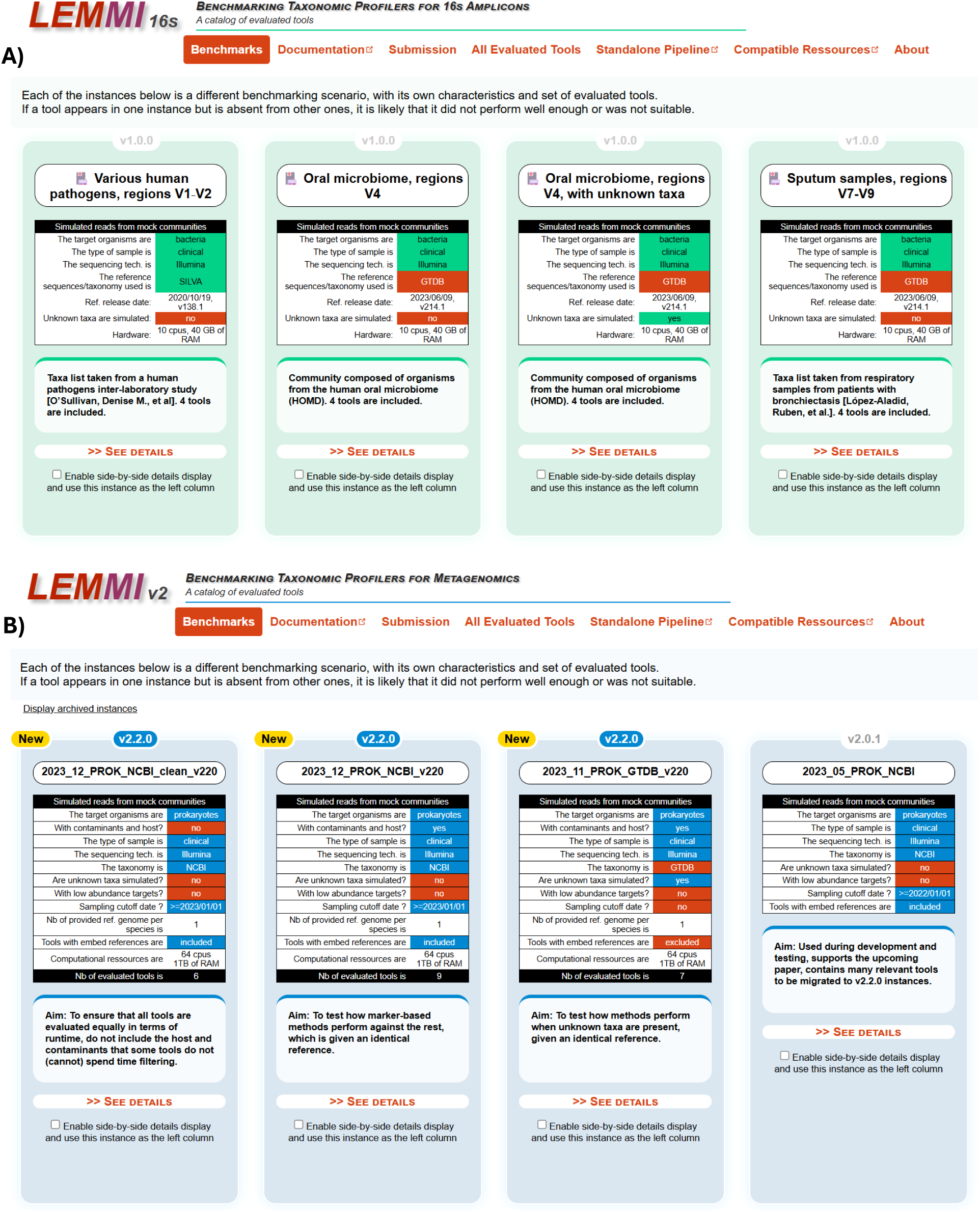
A) The home page of LEMMI16S. Different benchmark instances are presented with a brief description of theirs contents, highlighting key features such as the taxonomy used or the presence of unknown lineages. The version of LEMMI16S used to generate the benchmark is specified for each instance. B) Home page of LEMMIv2 for metagenomics with a similar layout.

A new feature of the LEMMIv2 benchmark is the inclusion of instances where host and contaminant taxa (non-targets) are present alongside the organisms of interest (targets). For example, reads representing the host genome should ideally not be included in predictions. This feature tests the tools’ ability to focus on relevant content, encouraging strategies, potentially involving the combination of tools into pipelines, to filter out or ignore irrelevant data before classification. In the LEMMI16S catalogue, users can explore instances simulating the amplification of various regions of the 16S gene from target organisms. Each LEMMIv2/16S instance encompasses multiple replicates representing different sampling of similar environments in the form of sequencing reads provided to candidate tools for generating predictions.

The online evaluation (Supplementary Figure 1) presents several widgets displaying various metrics summarizing prediction quality and resource consumption for each evaluated method. The top section presents averages across all samples, while subsequent sections allow users to explore the data sample by sample (Supplementary Figure 2). A widget consolidates key metrics for a user-selected shortlist of tools into a single spider plot (Figure 3). Another section of the website (“All Evaluated Tools”) offers useful information about each tool, including links to the code, relevant publications, and the container used for evaluation (Supplementary Figure 3).

**FIGURE 3.**
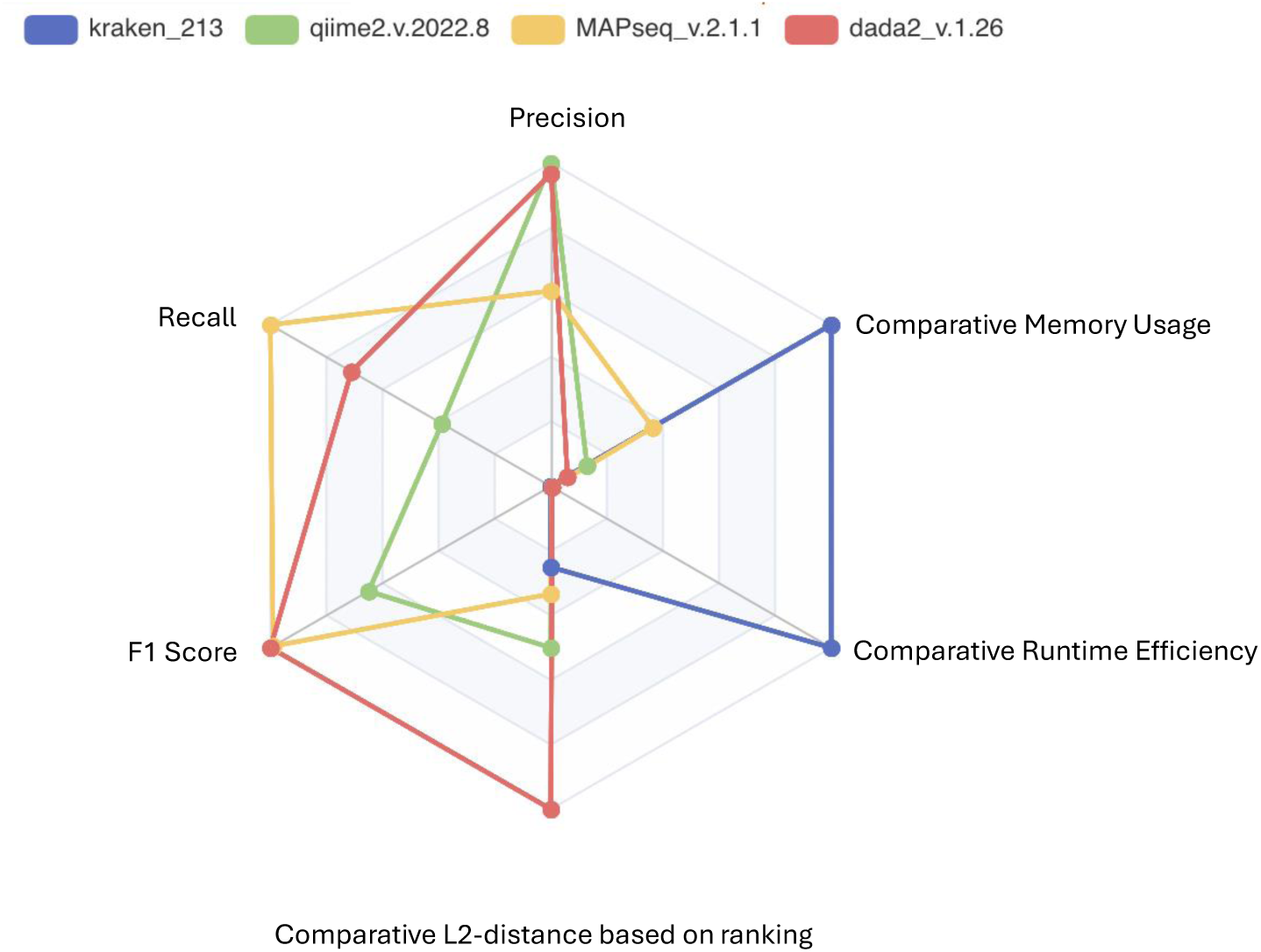
The LEMMI16S widget summarising key metrics. Users can select a shortlist of tools to display together.

#### The standalone pipeline

The standalone pipelines allow users to replicate the online benchmarks or create their own local instances with minimal effort. These pipelines can be easily deployed by cloning the code from the Git repositories (https://gitlab.com/ezlab/lemmi-v2/, https://gitlab.com/ezlab/lemmi16S/) and installing the required packages using the Conda installer, along with setting up a container engine (details provided below). Once set up, the LEMMIv2/16S pipeline automatically downloads and prepares the necessary reference repositories, generates benchmark instances, evaluates candidate tools, and displays the results on a web page. The only task left for the user is to complete the configuration files. Additionally, because LEMMIv2/16S are built on Snakemake (Köster & Rahmann, 2012), jobs can be efficiently distributed across CPU cores on a local machine or multiple nodes in a computing cluster.

##### Docker and Apptainer containers

LEMMIv2/16S ensures the long-term availability of exact tool versions for evaluation through the use of containers. A container functions as a self-contained evaluation unit, allowing the combination of multiple steps or even different methods within it. It receives input files from LEMMIv2/16S and generates the required outputs for evaluation. Resource usage, such as runtime and memory consumption, is measured to include all processes that occur when the container is deployed for a benchmarking task.

In LEMMIv1, Docker (https://www.docker.com/ last accessed 10.02.2015) was the only supported container engine. However, LEMMIv2/16S now supports both Docker and Apptainer. Apptainer (formerly Singularity) (Kurtzer et al., 2017) is widely adopted in HPC environments for its enhanced security features, avoiding some vulnerabilities found in Docker (Zhou et al., 2022), and its native integration with Snakemake (Köster & Rahmann, 2012). Additionally, Graphical Processing Units (GPUs), which are increasingly used in machine learning applications, can be utilised in LEMMIv2/16S with both container technologies, further expanding the platform’s capabilities.

All software containers that encapsulate candidate tools can either be sourced from online directories or accessed locally. Compatible containers, along with corresponding configuration files or datasets, are available at https://quay.io/user/ezlab/ and https://zenodo.org/communities/lembench/. Other repositories, such as https://hub.docker.com (last accessed 10.02.2025), can also be used to share and access compatible software containers, ensuring flexibility and broad accessibility for users to find the necessary resources for LEMMIv2/16S evaluations.

#### Sequences management

The LEMMIv2/16S pipeline manages microbial genomes for benchmarking through well-structured repositories. These repositories serve both as sources for generating *in-silico* reads and as references during tool evaluation.

##### Repositories

Support for GTDB is a new feature in LEMMIv2. Users can now build the LEMMIv2 sequence directory using prokaryotic, eukaryotic, and viral genome assemblies from GenBank, annotated with both NCBI taxonomy and GTDB metadata for prokaryotes. Each genome is validated to ensure it includes the necessary metadata; those lacking this information are discarded. LEMMIv2 ensures reproducibility by applying a user-defined cut-off date to filter the input material. Additionally, in this version, users can specify a host genome separately by providing a link to an assembly in the configuration, such as the human genome GRCh38.

Unlike LEMMIv1, LEMMIv2 does not require each genome to have a corresponding protein equivalent to be included in the repertoire. This previous requirement excluded many assemblies and limited LEMMIv1’s ability to represent the full diversity available in public databases. In LEMMIv2, control over the reference material during the benchmarking process is restricted to nucleotide-based approaches. However, tools that rely on protein references can still be evaluated by using a reference provided with the tool, similar to how marker gene-based tools are assessed, though this comes with some limitations (as detailed below).

In the case of LEMMI16S, the sequence repository is constructed using either the SILVA database (Pruesse et al., 2007) or a GTDB16S release (Parks et al., 2018) containing bacterial sequences. As with LEMMIv2, each amplicon is validated to ensure that the appropriate metadata is included. Filters are also applied to remove low-quality or redundant sequences, ensuring a well-defined reference repository.

##### Flexible control of the reference database

For all tools capable of creating a reference on demand, LEMMIv2/16S manages and provides the sequences that can be used as references (Figure 4A). In LEMMIv1, a genome exclusion approach was implemented: any genome used to generate sequencing reads was removed from the reference provided to the tool to avoid overfitting (Figure 4B). LEMMIv2 continues to utilise this approach.

**FIGURE 4.**
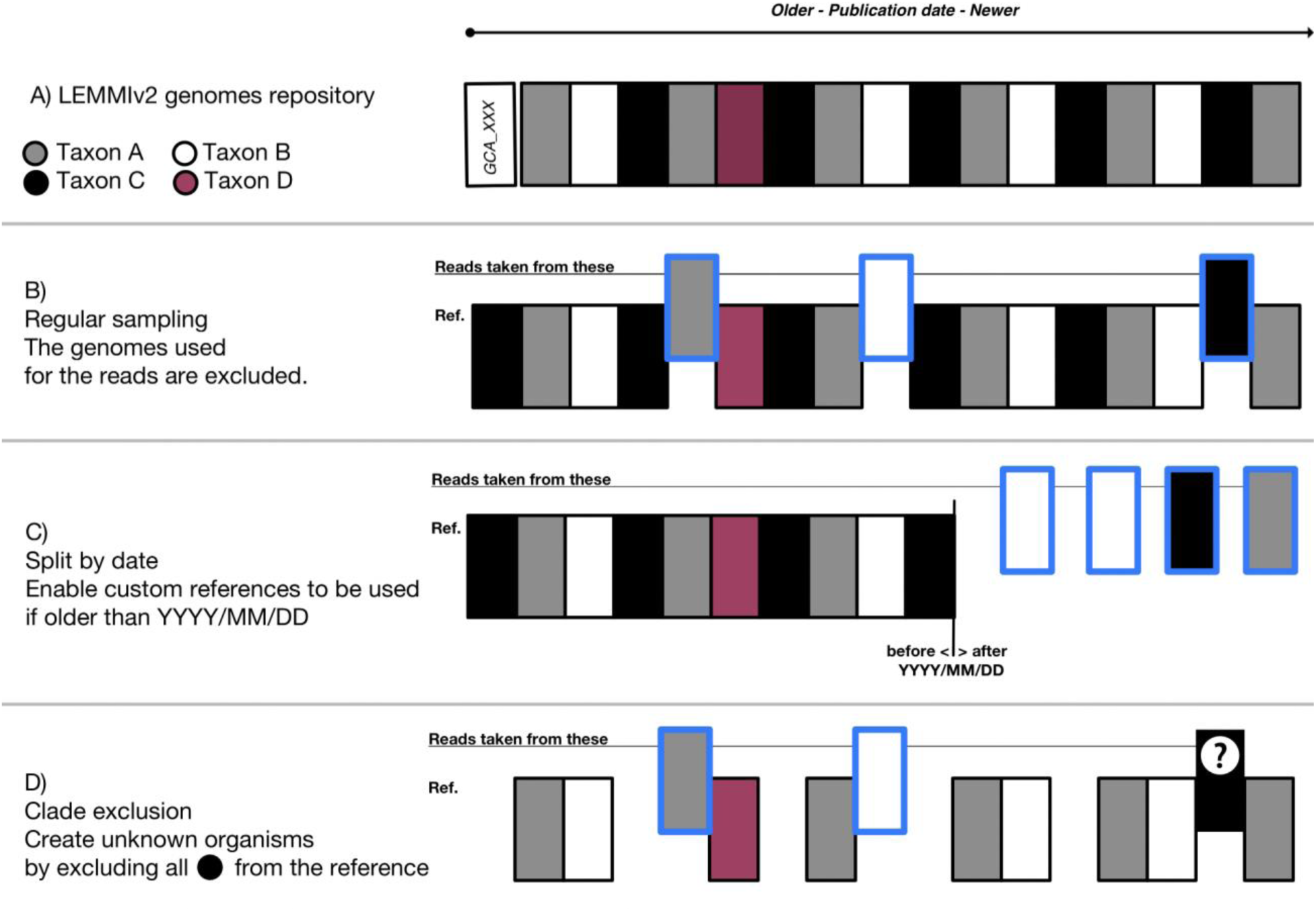
LEMMIv2 reads and references sampling. A) GenBank genome assemblies with sufficient metadata serve as starting material. Four species are represented by four colours (white, grey, black, purple). B) When creating an instance of LEMMIv2, genomes are randomly selected. At least two representatives are required for a species to be retained. These serve as the source of the reads, while the rest form the reference provided to the tools. C) If controlling the reference for all tools is not possible, (e.g. for tools based on curated marker genes), a cut-off date can be used to separate the reference from the reads. By defining a date that corresponds to the publication date of the most recent custom reference, LEMMIv2 will select only more recent genomes for the reads and keep all older genomes for the reference provided to the other tools. D) To increase complexity, entire species (in this case taxon C in black), are made unknown by removing all representatives from the reference provided to the tools.

However, this method is not applicable to tools that require an embedded reference, such as marker gene-based methods, where the reference database cannot be altered. Consequently, LEMMIv1 allowed all genomes to be kept as references during benchmarks including tools requiring embedded references, resulting in uniform overfitting bias by possibly including the exact source of the reads in the reference. This less-than-ideal strategy has been replaced in LEMMIv2. The new approach involves splitting the reference repository using a cut-off release date: genomes older than this date are used as references, while the more recent genomes are used to generate the reads (Figure 4C). This method allows tools with embedded references to be evaluated alongside other tools, provided that a cut-off date older than the release date of the embedded reference is set, ensuring that genomes used to generate the benchmarking reads were not available when the reference was created.

None of the tools evaluated with LEMMI16S require an embedded reference, so the cut-off strategy is not currently applicable, although it could be implemented in the future. The current version of LEMMI16S always returns the entire 16S marker as a reference, regardless of the specific region present in the reads. As with LEMMIv2, LEMMI16S excludes any amplicon used in the read-generation from the reference database for methods requiring model training.

##### Clade exclusion

In most samples, particularly those from new environmental niches, some taxa often lack close relatives in the reference database. This scenario can be effectively simulated using LEMMIv2/16S.

To provide a more realistic challenge, LEMMIv2/16S can designate unknown organisms, which helps distinguish tools that accurately handle reads from undiscovered taxa from those that generate numerous false positives in such contexts. To simulate the presence of unknown taxa using public data, LEMMIv2/16S allows users to designate certain lineages (at any taxonomic rank, such as species or family) as unknown during sample generation. Genomes or sequences from these designated lineages are excluded from the reference provided to the tools, regardless of their use in the read generation process (Figure 4D). Consequently, any reads belonging to these lineages must be reported as unknown to provide a correct prediction. It is important to note that this option should not be used when evaluating tools with embedded references, as the reference cannot be modified in those cases.

#### Minimum filter threshold to consider a taxon

Previous benchmarks have highlighted the importance of applying a minimum threshold of reads to validate the presence of a taxon with methods that classify all reads, in order to reduce false positives (McIntyre et al., 2017). This threshold may vary depending on the method used, but in real-life analyses and without calibration values, it can only be set arbitrarily and be the same of all methods. However, in a simulated environment, the known contents of the samples allow for the definition of an optimal threshold that maximises each tool’s performance. This threshold can then potentially be applied to real samples, provided that the simulation parameters (such as the number of reads, species richness, and abundance distribution) are representative enough of the actual sample.

The LEMMIv2/16S pipeline employs a strategy based on the aforementioned rationale to optimise the expected performance of each evaluated tool and provide guidance on setting an appropriate threshold for each. It achieves this by simulating two types of samples: “calibration samples” (indicated by the suffix c00x) and “evaluation samples” (indicated by the suffix e00x), using identical parameters. The taxa specified in the configuration file of the benchmarking instance are assigned random abundances according to the same distribution. While the dominant taxa may differ between the samples, the taxa present, their number, and the total number of reads remain consistent. The filtering threshold that maximises the quality of predictions in the calibration samples (by optimising the F1 score for taxa presence detection) is applied to the evaluation samples prior to calculating the metrics reported in the benchmark, ensuring that the results reflect the tool’s optimal performance. Additionally, the threshold for each tool is displayed in a dedicated widget, helping users understand calibration differences between tools.

### Comparison of simulation and real samples

To ensure realistic benchmarks, all LEMMIv2 and LEMMI16S instances are designed based on actual biological samples. To assess how well the simulation aligns with the real-world sample, we compared specific features of both datasets for a given instance. For example, a dental plaque sample from a healthy patient was used to guide the simulation process for LEMMIv2. The parameters for this pipeline were estimated from an initial analysis of the sample, reported in the LEMMIv2 configuration file https://lemmi.ezlab.org/data/2023_05_PROK_NCBI.yaml. The number of reads, read lengths, distributions, compositions (e.g., number of organisms and taxa), sequencing technology and related settings were carefully tailored to ensure that the resulting LEMMIv2 instance (https://lemmi.ezlab.org/2023_05_PROK_NCBI) accurately mirrored the characteristics of the original dental plaque sample.

A similar method was employed for LEMMI16S, where parameters for the simulation were derived from samples collected from the respiratory tract, specifically the dataset NCBI SRA reference SRR22879376.

Supplementary Table 1 and Supplementary Figure 4 illustrate the similarities between the simulated and real samples in terms of reads composition, species distribution, and GC content. These results demonstrate the ability of the instance generation process to faithfully replicate key characteristics of the original biological material.

### The evaluation results

#### LEMMIv2 for metagenomics

LEMMIv1 was limited to generating samples representing Illumina’s short read technology. To expand the scope of tool evaluations, particularly for those dedicated to long-read analysis, LEMMIv2 now includes the ability to simulate Oxford Nanopore Technology data. This enhancement allows the evaluation of tools specifically designed for long-read applications, as well as the assessment of methods originally developed for short-read inputs when they are applied to long-read datasets.

The prokaryotic profilers primarily designed for short reads that have been evaluated at the time of writing include MetaPhlAn v4.0.2 (Blanco-Míguez et al., 2023) and v3.0.13 (Beghini et al., 2021), mOTUs v3.0.1 (Ruscheweyh et al., 2022), Sourmash v4.2.3 (Brown & Irber, 2016), KMCP v0.9.0 (Shen et al., 2023), Kraken v2.1.2 (Wood et al., 2019), Centrifuge v1.0.4 (Kim et al., 2016), Ganon v2.0.0 (Piro & Reinert, 2023) and v1.1.0 (Piro et al., 2020), CCMetagen v1.4.0 (Marcelino et al., 2020), Metacache v2.2.0 (Müller et al., 2017), Centrifuger v1.0.0 (Song & Langmead, 2024), and Metabuli v1.0.2 (Kim & Steinegger, 2024). Where applicable, these tools were also evaluated on long reads and viral datasets. The viral prediction pipeline VirMet v1.0.0 (https://github.com/medvir/VirMet last accessed 10.02.2025) was evaluated only on viral datasets. Metamaps v0.1.0 (Dilthey et al., 2019) and MMseqs v2.12 (Steinegger & Söding, 2017) were assessed using Oxford Nanopore Technology simulations.

Certain tools were considered for evaluation but did not scale efficiently in terms of runtime under the conditions tested; LEMMI reports these entries as they are benchmarking results. These include kASA (Weging et al., 2021) and MTSv (Furstenau et al., 2022), which are mentioned on the LEMMI website but not yet evaluated due to current performance limitations. Deepmicrobes (Liang et al., 2020) was evaluated but its predictions were limited to the genus level and above.

##### Clinical samples

Dental plaque samples were analysed using several candidate tools to identify approximately fifty bacterial species per sample (https://lemmi.ezlab.org/2023_12_PROK_NCBI_v220). These samples simulated Illumina short reads and were classified under the NCBI taxonomy. The simulated mix also included human host and fungal reads as non-target organisms. To avoid overfitting, the repository was split into reference and sample using a cut-off date set later than the release of MetaPhlAn 4.0.2, the tool with the most recent embedded reference (as depicted in Figure 4C). All methods performing read-level classification were given the opportunity to filter out human host and fungal reads before analysing the remaining sequences to classify the target organisms (Supplementary Table 2).

At the species level, marker-based tools delivered the most accurate predictions of taxa presence, despite their inability to remove host and contaminant reads prior to analysis. MetaPhlAn 4 emerged as the top performer in terms of F1 score, with the highest precision and recall (Supplementary Figure 5). Among k-mer-based methods processing the LEMMI-provided reference, Kraken, Ganon, Sourmash, and KMCP showed similar performance levels. To manage resource limits (memory, disk space, and runtime), each species in the reference included only one representative genome. However, some tools capable of handling up to five reference genomes per species were re-evaluated with a more comprehensive reference, and these results were marked as (toolname)_all. Interestingly, this led to a negative impact on recall for some tools like Kraken, as previously observed by (Nasko et al., 2018).

Centrifuge showed a slightly lower F1 score than the top performers, while Metabuli and Centrifuger ranked lowest due to low precision values. LEMMIv2 automatically applied a filtering threshold derived from calibration samples, optimised to maximise the F1 score at the species level; the values are reported in Table 1. Regarding the L2 distance to the truth (measuring the distance between predicted and actual abundances), KCMP performed the best, followed by Kraken and mOTUs, while MetaPhlAn 4 had the least accurate abundance estimates (Supplementary Table 3).

**Table 1.** Minimal number of reads required to consider a taxon as being present at the *species* level in the evaluation samples of the instance 2023_12_PROK_NCBI_v220. This lower detection limit maximizes the F1-score on the calibration samples and is applied to the evaluation samples.

| Tool name | Value |
| --- | --- |
| metaphlan_402 | 0 |
| motus_301 | 0 |
| kraken_212 | 1530 |
| kraken_212_all | 2548 |
| ganon_200 | 1054 |
| sourmash_423_all | 988 |
| ganon_200_all | 1126 |
| kmcp_090 | 2722 |
| sourmash_423 | 1141 |
| centrifuge_104 | 4341 |
| metabuli_102 | 5638 |
| centrifuger_100 | 5190 |

At higher taxonomic levels (genus and above), the performance gap between MetaPhlAn 4 and the other tools narrowed. mOTUs improved its precision significantly, becoming the best performer in terms of F1 score at the genus level (Supplementary Figure 6).

Additional tools such as Metacache, CCMetagen, and Deepmicrobes were evaluated alongside many of the previously mentioned tools in an earlier LEMMI instance, which is available at https://lemmi.ezlab.org/2023_05_PROK_NCBI. This instance represents the same dental plaque sample type. These tools ranked lower in F1 scores for species detection and were not reevaluated on the latest instance.

##### Runtime and memory evaluation

To fairly assess runtime without penalizing tools that first filter out host and contaminant reads, compared to those that do not, a clean version of the previous LEMMI instance containing only target bacterial reads was created (https://lemmi.ezlab.org/2023_12_PROK_NCBI_clean_v220). This benchmark revealed that Ganon was the fastest tool, processing 3 million Illumina 150 base pair paired-end reads in just half a minute. On the other hand, MetaPhlAn 4 and KMCP were the slowest, each taking approximately five minutes to complete the analysis (Supplementary Figure 7A).

In terms of memory usage, Sourmash and mOTUs stood out as the most efficient tools (Supplementary Figure 7B). Interestingly, when evaluating tools with one representative genome per species versus five genomes per species, Sourmash maintained its memory footprint, even with a more comprehensive reference (e.g., sourmash_423 vs. sourmash_423_all on https://lemmi.ezlab.org/2023_12_PROK_NCBI_v220). This contrasts with other tools, which generally increased their memory requirements when using more extensive references.

##### Unknown taxa

The benchmarking instance presented above only includes taxa that are currently represented in the database, making it easier to solve than real-world samples. To address this, a similar instance was created (https://lemmi.ezlab.org/2023_11_PROK_GTDB_v220), where one in every three taxa was marked as unknown (as shown in Figure 4D). This instance is suitable only for evaluating tools that can construct their reference from the provided material, and the target taxonomy is set to GTDB.

In this more challenging scenario, Sourmash achieved the best results, outperforming other top tools such as Kraken, KMCP, and Ganon by maintaining high recall and precision, leading to the highest F1 score. KMCP was notably affected by the inclusion of unknown taxa, showing a drop in precision at the species level (Supplementary Figure 7C). Despite this, KMCP still minimized the L2 error in abundance estimation. This is because its false predictions tended to involve low-abundance taxa, while highly abundant taxa were accurately reported.

##### Environmental samples

A benchmarking instance (https://lemmi.ezlab.org/2022_03_PROK_NCBI_2) was created based on a study of a meromictic lake (Saini et al., 2022). The aim was to estimate the prokaryotic fraction, consisting of about 20 species at the species rank. However, all tools tested showed poor precision and recall (Supplementary Figure 7D). The poor recall could be attributed to the lack of reference genomes for niche environmental taxa in tools with embedded references, such as MetaPhlAn and mOTUs. But designing a LEMMI instance with a reference genome for every species should have enabled other tools to perform similarly well as they did with human samples. A likely explanation for the poor results is either the greater genomic distance between species in this environmental instance and their reference compared to clinical samples or an improper labelling of some reference genomes under the NCBI taxonomy. This mismatch between the NCBI taxonomy and the true genomic distances between the organisms placed in the reads and in the reference may have prevented tools from correctly associating sequences with the appropriate taxonomic identifiers more frequently than with human microbiome simulations.

To test this hypothesis, two instances with identical parameters were created (https://lemmi.ezlab.org/2024_01_PROK_NCBI_v220 and https://lemmi.ezlab.org/2024_01_PROK_GTDB_v220), differing only in the taxonomic system used (NCBI vs. GTDB). GTDB is a taxonomic system constructed with genomic distances. When tested with KMCP, the F1-score at the species level was significantly higher for the GTDB taxonomy instance (0.8) compared to the NCBI taxonomy instance (0.34). This improvement was also reflected in abundance estimation, where the L2 distance dropped from 0.6 (NCBI taxonomy) to 0.03 (GTDB taxonomy) for sample e001, highlighting the impact of the taxonomic representation based on genomic distances when analysing understudied niche compared to well characterised and highly sampled human microbiomes.

We also generated an instance inspired by the same study (Saini et al., 2022), focusing on the eukaryote fraction of the sample (https://lemmi.ezlab.org/2022_01_EUK_NCBI). In this case, Sourmash demonstrated the best performance in species identification, recovering an average of 3.5 out of 4 species while maintaining a much lower false prediction rate than other high-recall tools (Supplementary Figure 7E). Marker-based tools were not included in this instance.

##### Viral samples

Samples containing eight low-abundance viruses (approximately 300 reads each among 5 million bacterial reads) were generated to simulate conditions similar to pathogen detection (https://lemmi.ezlab.org/2021_9_VIR_NCBI). In this scenario, Sourmash performed poorly (Supplementary Figure 7F), while Kraken outperformed all other tools, including VirMet, which was the only dedicated viral pipeline evaluated. Kraken achieved the best F1 score at the species level, with VirMet ranking second.

##### Long reads

LEMMIv2 also supports the simulation of Oxford Nanopore long reads. The clinical samples discussed earlier were generated in an additional LEMMI instance (https://lemmi.ezlab.org/2022_04_PROK_NCBI_LR2) consisting of 20,000 reads with lengths ranging from 57 to 32,389 base pairs, averaging 1,096 base pairs, and covering approximately 70 bacterial species. Several tools originally designed for short reads were evaluated, and only those that successfully completed the analysis with meaningful results were reported. Centrifuge demonstrated solid performance, though it did not match Metamaps in terms of recall and precision at the species level (Supplementary Figure 7G). However, Centrifuge was significantly faster, completing the analysis hundreds of times quicker.

#### LEMMI16S

LEMMI16S currently focuses on Illumina short reads, with long reads considered for future benchmarks. In the current version, four candidate tools have been included: QIIME 2 (version v2022.8) (Bolyen et al., 2019), DADA2 (version v1.26) (Callahan et al., 2016), Kraken2 (version v2.13) (Wood et al., 2019), and MAPseq (version v2.1.1) (Matias Rodrigues et al., 2017). These tools were subjected to comprehensive evaluation under different scenarios, with simulated reads being generated from various amplicon repositories (i.e., SILVA release_138.1 and GTDB release 214.0), hypervariable regions (i.e., V1-V2, V4, and V7-V9), and bacterial compositions. Other tools, including mothur (version v1.47) (Schloss et al., 2009) and SPINGO (version v1.3) (Allard et al., 2015), were also considered for evaluation, but their runtime and prediction accuracy fell below expectations, requiring further investigation before public release.

The next section describes the instances in which these candidate tools were evaluated. Note that alternative 16S rRNA databases, such as GreenGenes (DeSantis et al., 2006) and RDP (Cole et al., 2009), were not considered, primarily because they have not seen updates in recent years. Analytic methods requiring complete amplicons or hypervariable regions exceeding 400 bp were excluded from consideration primarily as a result of constraints imposed by short-read sequencing technology, in which read length is restricted to 250 bp for both forward and reverse directions, and several tools mandate a minimum overlap between these read pairs. Furthermore, the analysis at the species level was omitted, as it has been suggested that species assignment for short 16S sequences necessitates 100% identity and distinguishing between species is inherently challenging due to the identical or highly homologous nature of 16S rRNA genes among certain distinct species (Edgar, 2018).

##### Human pathogens samples

The dataset was inspired by the inter-laboratory study proposed by O’Sullivan et al. (O’Sullivan et al., 2021). It contains fifteen popular human pathogens species, with their abundance being randomly represented in five instances (3 for calibration and 2 for evaluation). The species amplicons were sourced from the SILVA repository. For each instance, 845,000 reads were generated by LEMMI16S (length=250bp, mean=400bp, std=10bp) from the V1V2 region. All candidate tools were evaluated with this dataset, considering their default stages and parameters (https://lemmi16S.ezlab.org/alfa_v1v2_SILVA_gcn, https://zenodo.org/records/8343988). Predictions at the family and genus levels were reported, as they indicated the most variations in the results. At these levels, all candidate tools were found to perform suitably, with subtle differences between all the metrics. QIIME 2_v2022.8 achieved the highest F1 score, DADA2_v1.26 excelled abundance estimation (with the lowest L2 distance to truth), and Kraken2_v213 was the fastest and most memory-efficient among the evaluated tools.

##### Human Oral Microbiome samples with unknown organisms

This dataset includes 74 genera from the Human Oral Microbiome Database (eHOMD) (Chen et al., 2010) are represented. The LEMMI16S workflow was configured to utilize the GTDB SSU database and generate five samples (3 for calibration and 2 for evaluation), each with 500,000 reads (length=250bp, mean=400bp, std=10bp) from the V4 region (a characteristic region used for studying HOM composition (Dashper et al., 2019)). Furthermore, an additional sample was created by randomly designating 14 organisms as unknown while retaining the other parameters (available at https://lemmi16S.ezlab.org/HOMD_v4_GTDB_gcn, https://lemmi16S.ezlab.org/HOMD_v4_unknown_GTDB_gcn, and https://zenodo.org/records/8344037). Supplementary Figure 8 illustrates that this condition slightly impacts the performance of the tools, with DADA2_v1.26 leading in F1 score and L2 distance in both scenarios, while Kraken excels in computational speed and memory efficiency.

##### Respiratory samples

This dataset comprises a taxa list derived from the experiment conducted on respiratory samples from patients with bronchiectasis by López-Aladid et al (López-Aladid et al., 2023). The samples (3 for calibration and 2 for evaluation) were subjected to the generation of 500,000 reads (length = 250bp, mean = 400bp, std = 10bp) from the V7–V9 region, representing 20 genera with varying abundance levels. The amplicons were sourced from the GTDB repository. At the Family and genus levels, low performance is demonstrated by DADA2_v1.26, while MAPseq_v2.1.1 exhibits clear superiority in terms of the F1-score. Remarkably, QIIME 2_v2022.8 showcases the best abundance estimation. In contrast, Kraken stands out as the fastest and the least RAM-intensive tool but performs poorly in other metrics, including F1-score and L2 distance.

## Discussion

Metagenomics practitioners, both users and developers, will find LEMMIv2/16S to be a versatile framework that enables the simulation of sequencing reads corresponding to samples they are working on or interested in. This functionality allows users to easily evaluate existing tools compatible with LEMMI, including their own developments, gaining insights into which features influence a tool’s behaviour, runtime, and resource consumption. One such feature is the proportion of unknown lineages present in the sample being analysed. Our benchmarks, based on public datasets, enable the rapid creation of new data, and the ability to selectively exclude specific genomes from the reference, which allows for accurate simulation of the challenges posed by the incomplete sampling of microbial diversity.

In addition, the benchmark maintained by the LEMMI team at https://lemmi.ezlab.org and https://lemmi16S.ezlab.org simplifies the process of centralising performance data for these tools, allowing for a quickly updated and widely shared list of evaluated methods. These platforms offer the community an ongoing and incremental evaluation resource. As of this writing, the benchmarks have included both widely popular and well-established tools, as well as newer, less-known options. Many tools are still awaiting evaluation, and we encourage developers to utilise LEMMI and submit their containers for public benchmarking.

When designing an experiment to study a microbial community, it’s essential to consider the classification algorithm and reference database independently to optimize their fit to the target material. A poorly chosen database can undermine the validity of the results (Gihawi et al., 2023). If resources and time permit, selecting a custom genome set is recommended rather than relying on default options, unless a comprehensive third-party database like https://benlangmead.github.io/aws-indexes/k2 (last accessed 10.02.2025) is preferred. In all cases, to accurately evaluate a method’s predictive power, the evaluation should be independent of any prepackaged reference constructed at an arbitrary date.

In line with this principle, LEMMIv2/16S ensures that, where possible, all tools are evaluated with the same reference material. This approach can reduce performance gaps between tools with similar methodologies, possibly indicating that transitioning to a new method may be unnecessary unless specific dataset features justify it (e.g., Supplementary Figure 5 shows Kraken, Ganon, KMCP, and Sourmash producing comparable results, while Supplementary Figure 7C demonstrates Sourmash’s superior performance when unknown organisms are present). This setup does not oversell minor progress but ensures that significant advancement in classification capabilities by a new tool will be clearly apparent through LEMMIv2/16S’s benchmarks in the future.

The evaluation of tools with embedded references (e.g., marker-based methods such as those included in this report) is essential for offering a comprehensive view of available options. To avoid an overfit of the benchmark, a cut-off date is used to ensure that the reads do not directly match the genomes likely used to build the markers. However, with the LEMMI approach, these tools cannot be fully tested in scenarios where the presence of unknown organisms simulates more realistic samples.

The work invested in the selection of marker genes for tools like MetaPhlAn and mOTUs undoubtedly results in high accuracy, with much of their strength tied to their reference datasets. It is important for LEMMI users to recognize that comparisons between tools with embedded references and those using controlled, external references serve different purposes. While the former demonstrates the effectiveness of a tool’s internal reference set, the latter emphasises the tool’s general classification performance. Therefore, when assessing tools, users should differentiate between these two approaches to make informed decisions based on their specific research needs.

In this paper, we emphasise that using the GTDB taxonomy to structure the reference database significantly enhances the analysis of niche environments compared to well-characterised human-related communities. For researchers working with these less-characterised environments, it is advisable to choose methods that can directly handle GTDB taxonomy or utilise tools that can convert GTDB data into an NCBI-like taxonomy (e.g., using https://github.com/nick-youngblut/gtdb_to_taxdump last accessed 10.02.2025). Additionally, marker-based methods should ensure compatibility with the GTDB framework to maximise their effectiveness in diverse and novel environments.

In addition to considering the time and memory required for sample analysis, the time required to process a reference may also be a criterion if frequent updates are needed. LEMMIv2/16S reports this information. Note that some tools, such as Ganon, can update their reference without reprocessing the existing part, so users will need to judge whether the initial construction as reported on LEMMI and the update process are different. We have found that increasing the size of the reference, for example, from one representative genome per species to five representatives, affects memory usage during sample analysis in different ways across tools. A tool performing well with 10,000 genomes may not scale effectively and remain the best when the reference expands to 50,000 genomes.

One might wonder whether deep learning approaches will bring significant advances or whether a plateau in classification performance has been reached. In this work, we evaluated Deepmicrobes with LEMMIv2, but its performance is not yet satisfactory. The containerization approach used by the pipeline allows for the use of GPUs in both reference construction and sample analysis, which will certainly cover future methods to be evaluated.

All tools evaluated in LEMMIv2/16S come with a public container, ensuring they can operate in a standardised environment (e.g., UNIX machines with dozens of CPUs, hundreds of GB of RAM, and terabytes of storage). The mandatory use of containers enhances reproducibility and simplifies the deployment of the benchmark. However, some software may require specific hardware or libraries for optimal performance, and portability cannot always be guaranteed without rebuilding the containers from source. Some tools have claimed high performance on desktop hardware (e.g., laptops) and LEMMIv2/16S can be used on such machines to test these claims against realistically sized datasets. As of now, we have not identified any examples in our benchmarks that behave well in these conditions.

Long read technologies represent a significant advancement in sample profiling, as they capture more information within each read and reduce the risk of classification errors caused by short, indistinguishable sequences. LEMMIv2’s simulation of Oxford Nanopore technologies has shown that MetaMaps, a method specifically designed for long reads, outperforms Centrifuge, which was originally developed for short reads but still managed to provide valuable results.

Future LEMMIv2 developments should include simulations for PacBio. Additionally, the LEMMI16S platform could benefit from integrating long read simulations, as long reads are increasingly relevant in the field of amplicon sequencing (Buetas et al., 2024).

LEMMIv1 included evaluations of read binning because tools like Kraken were initially designed for read classification with companion scripts for profiling. However, to maintain a focus on profiling in LEMMIv2, read binning aspects are no longer evaluated, as not all tools support read-level analysis. A future platform dedicated to read binning, leveraging the existing software containers, could be established to address these tools and their specific needs.

### Conclusions

In this report, we introduced the latest version of LEMMI: LEMMIv2 for metagenomics and its sister platform, LEMMI16S, for evaluating amplicon sequencing methods. We detailed their key features and shared some of the results obtained, which were used to build tool catalogues on https://lemmi.ezlab.org and https://lemmi16S.ezlab.org. We demonstrate how LEMMI enables re-evaluation of established methods and new software releases in unbiased metagenomics scenarios within an independent computational environment. The goal of LEMMI is to maintain an up-to-date catalogue and encourage both tool users and developers to visit these websites, access the standalone pipelines, design their own benchmarks, and submit their developments for public evaluation. This collaborative effort will contribute to a comprehensive knowledge base on metagenomics and 16S amplicon sequencing methods.

## Methods

The LEMMIv2/16S standalone pipelines are built around several Snakemake (Köster & Rahmann, 2012) workflows. The features not specifically attributed to one of the pipelines in this section are common to both. Both have parameters defined in different YAML configuration files for the global configuration (e.g. https://gitlab.com/ezlab/lemmi-v2/-/raw/v2.2.0/config/config.yaml.default), the creation of instances (e.g. https://gitlab.com/ezlab/lemmi-v2/-/raw/v2.2.0/benchmark/yaml/instances/demo_prok_gtdb.yaml) and the execution of candidate methods (e.g. https://gitlab.com/ezlab/lemmi-v2/-/raw/v2.2.0/benchmark/yaml/runs/kraken_212.demo.yaml).

### Containerization

Several pre- and post-processing tasks in the pipeline, as well as all evaluated methods, are wrapped in software containers and run either on Singularity (Kurtzer et al., 2017) through its native integration with Snakemake, or on Docker (https://www.docker.com last accessed 10.02.2025) through a custom integration with Snakemake. The resources used by the candidate methods are reported by tracking the time the container is loaded and the peak RSS memory reached by any process running inside the container.

### Repository construction

LEMMIv2 is based on the genbank list of assemblies which is obtained from https://ftp.ncbi.nlm.nih.gov/genomes/ASSEMBLY_REPORTS/assembly_summary_genbank.txt (last accessed 01.11.2023). It can be filtered to remove entries older and newer than the date range specified in the configuration file. The results presented in this manuscript are based on all genomes older than 2023/11/01. Each species lineage is validated using the ETE Toolkit 3 library (Huerta-Cepas et al., 2016) set with NCBI dmp files corresponding to a user-defined date. The results presented in this manuscript use 2023/11/01. Entries without a valid lineage and without an FTP URL are dropped. In addition, the metadata corresponding to a release of the GTDB taxonomy is downloaded and parsed, and the genomes present in that taxonomy are annotated with extra information. The results presented in this manuscript are based on the GTDB release 207. All remaining genomes matching the lineages specified in the config file are then downloaded up to a user-defined limit of genomes per species (results in this manuscript: five genomes per species covering all Bacteria and Archaea as well as microbial eukaryotes including all Fungi) to form the LEMMIv2 repository. An equivalent of the NCBI dmp files are generated for GTDB using https://github.com/nick-youngblut/gtdb_to_taxdump (last accessed 10.02.2025). A host species is also provided. For the work presented in this manuscript, the human genome version GRCh38 was used.

In the case of LEMMI16S, the sequence repository is built using either the SILVA database (Glöckner et al., 2017) or a GTDB16S release (Parks et al., 2018), both of which contain bacterial 16S sequences. Poor-quality, redundant, and short sequences are removed from the final repository using the RESCRIPt tool and tutorial (Robeson et al., 2021). The resultant sequences and their taxonomy are conserved for future stages. In this case, all sequences in the repository always represent the complete 16S genes. The hyper-variable regions to be amplified are defined and extracted from the repository sequences during the generation of the LEMMI16S instance.

### Instance creation

The LEMMIv2 configuration file defining an instance is parsed to obtain the target clade (prokaryotes, eukaryotes or viruses) and the contaminant clades (prokaryotes, eukaryotes, and/or viruses). The relative proportion of each clade plus the host is defined, and the number of genomes to use for each clade is also taken from the configuration file. If specified, the genomes are filtered to match the specific lineages before the expected number of genomes are sampled from the repository. For each clade (prokaryotes, eukaryotes, viruses), a lognormal distribution is used to randomly define the individual contribution of each genome to the relative abundance. This sampling operation is performed for each sample defined in the configuration file, except for those labelled as negative samples which are generated without the target clade. The list of genomes remaining as a reference is created by excluding those selected in the previous step, as well as all genomes belonging to the lineages marked as “unknown” in the configuration file. If a cut-off date is defined for selecting the reference, the genomes for creating the reads are sampled within the genomes more recent than this date, and the reference is made up of all genomes older than the cut-off date. For Illumina sequencing, the ART reads simulator (Huang et al., 2012) is used to generate the samples via the readSimulator library (https://github.com/wanyuac/readSimulator last accessed 10.02.2025); several parameters such as the number of reads or the Illumina sequencing chemistry need to be specified. If the sequencing technology is Nanopore, NanoSim (Yang et al., 2017) is used.

The LEMMI16S pipeline applies two filters to select amplicons and produce simulated reads. A taxon filter allows the selection of amplicons from a specific lineage, and a region filter enables the conservation of only those that contain the particular zone in the 16S gene that is mentioned in the instance config file. The resultant amplicons, which come from the specific taxa and include the indicated region, are split into two datasets (query and reference). The reference dataset is reserved for building the model reference. The query dataset is used as the input for the simulator tool, which produces the ‘synthetic’ reads. The instance configuration file allows setting the sample composition (organisms or taxonomic rank), the target 16S regions (e.g. V1-V2), the query/reference percentage, the total number of reads to be generated, and the simulator properties. If a particular taxonomic rank is marked as unknown, all its sequences are removed from the reference dataset but retained in the query dataset.

### Reference processing by the candidate methods

Before running an evaluation, the LEMMIv2 pipeline places all genomic and taxonomic files needed to build the references for the selected instance into a dedicated temporary folder, organised into three sections: the host genome, reference genomes for target organisms, and reference genomes for unwanted organisms to be excluded from the report. The candidate container is loaded and the scripts LEMMI_process_ref_host.sh, LEMMI_process_ref_targets.sh and LEMMI_process_ref_contaminants.sh are called in sequence. The candidate method container must produce three databases compatible with its classification procedure for the next step. Methods that cannot fulfil some or all of these tasks will instead execute a sleep command for 30 seconds to acknowledge the completion of this step.

LEMMI16S replaces the LEMMI_process scripts with the LEMMI16S_training.sh script. It contains the candidate method’s instructions to train a machine learning model or build a reference database (according to the method strategy) from the reference dataset. Similar to LEMMIv2, a sleep command for 30 seconds is executed when no reference needs to be built.

### Analysis of samples by the candidate methods

To obtain the predictions necessary to evaluate a candidate tool, the LEMMIv2 pipeline places all *in-silico* reads in the dedicated temporary folder containing the references built in the previous steps. It then loads the candidate container and invokes the LEMMI_analysis.sh script. The candidate method container must respond to this call by providing a taxonomic profile of the sample based on the references available in the folder, unless the method comes with an embedded reference and did not create any at the previous step. If applicable, the method will first filter out the host reads, then filter out the contaminant reads, and then classify the remaining reads to produce a report with four columns: organism name (binomial for species, name for the parent clades), taxid, number of reads, and relative abundance of genome copies; the abundance includes the unclassified fraction. The reported parent taxonomic ranks may or may not include the sum of the child ranks, and LEMMIv2 will calculate the missing values depending on the option selected.

Similarly, LEMMI16S invokes the LEMMI16S_analysis.sh script to copy the *in-silico* reads generated from the query dataset into a temporary folder. It then uses the reference model to classify them. If necessary, the method will call auxiliary tools to perform preliminary tasks before generating the final classification. The four-columns format presented above is used to produce the final report. For the tools that report OTU or ASV classification, the number of reads is computed by consulting the size of the OTU or ASV.

### Evaluation of the predictions

The predictions provided by the candidate methods are summed and filtered to ensure that they contain only entries consistent with the taxonomic system used by LEMMI and provided during the reference construction step. The samples labelled ‘calibration’ are then used to find the filtering threshold that maximises the F1 score for taxa presence/absence predictions at each taxonomic rank considered (the results presented in this manuscript cover phylum, class, order, family, genus and species). This method-specific threshold is then applied to the taxonomic profiles reported for all samples labelled ‘Evaluation’. The following metrics are calculated from the filtered taxonomic profiles: precision in taxa presence/absence predictions, recall in taxa presence/absence predictions, F1 score in taxa presence/absence predictions, and L2 error in relative abundance estimation of genome copy number. The numbers of true positives and false positives in taxa presence/absence predictions are also reported. Bray-Curtis dissimilarity is used to create a heatmap of the dissimilarity of each tool’s predictions.

### Web app to explore the results

A web application presents all the results, including the prediction accuracy metrics mentioned above, as well as the runtime and RSS memory peak of the different tasks performed by the candidate methods. The web app is built with Vue.js (https://vuejs.org/ last accessed 30.06.2024), deployed using npm (https://www.npmjs.com/ last accessed 30.06.2024), and uses Apache ECharts (https://echarts.apache.org/ last accessed 30.06.2024) to generate the plots. The web app is packed in the LEMMI master container, which is also required to perform other tasks throughout the benchmarking lifecycle.

### Companion scripts

Several candidate methods natively report read abundance rather than genome copy abundance. To make all methods comparable in this respect, a post-processing script, identical for all methods, was added to candidate containers where necessary to normalise the profile of reads to a profile of genome copies, using the information provided by LEMMIv2 to the method to build the reference.

### Computational resources

The results presented in this manuscript were produced using Singularity version 3.8.1-1.el8 running on a dedicated server with 48 Intel® Xeon® w5-3425 CPUs, 500 GB DDR5 RAM, and 250 TB disk storage.

## Acknowledgements

This research was funded in part by the Swiss National Science Foundation (SNSF) 310030_189062 to EZ. AB acknowledges support from the Federal Commission for Scholarships for Foreign Students for the Swiss Government Excellence Scholarship (ESKAS No. 2022.0531).

## Author contributions

MS, AB, and EZ conceived the study. MS and AB coded the pipeline. MS and MB coded the webapp. MS, AB, and MM conducted the analyses. EZ led the project. All authors wrote the manuscript.

## Supplementary Material

**Supplementary Figure 1.**
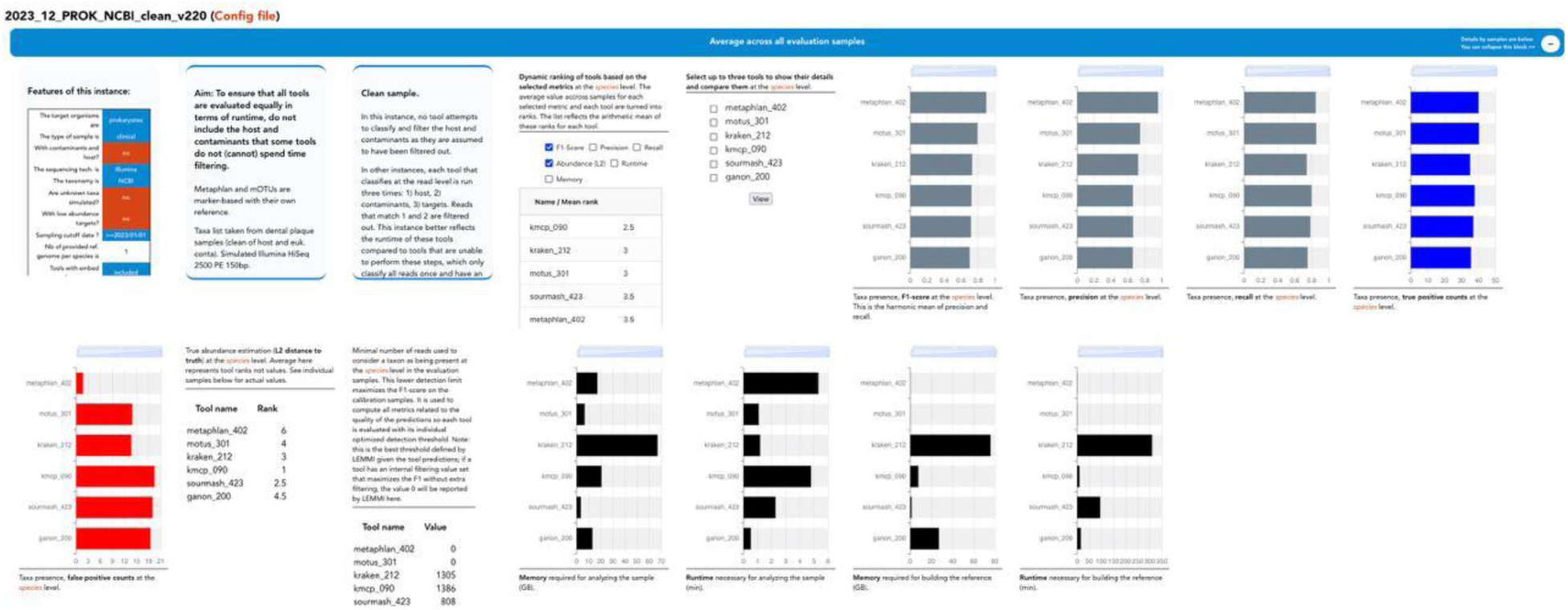
Detail page of a LEMMIv2 instance, showing different widgets reporting average values for all the samples or metrics that are not specific to a sample such as resources for building the reference.

**Supplementary Figure 2.**
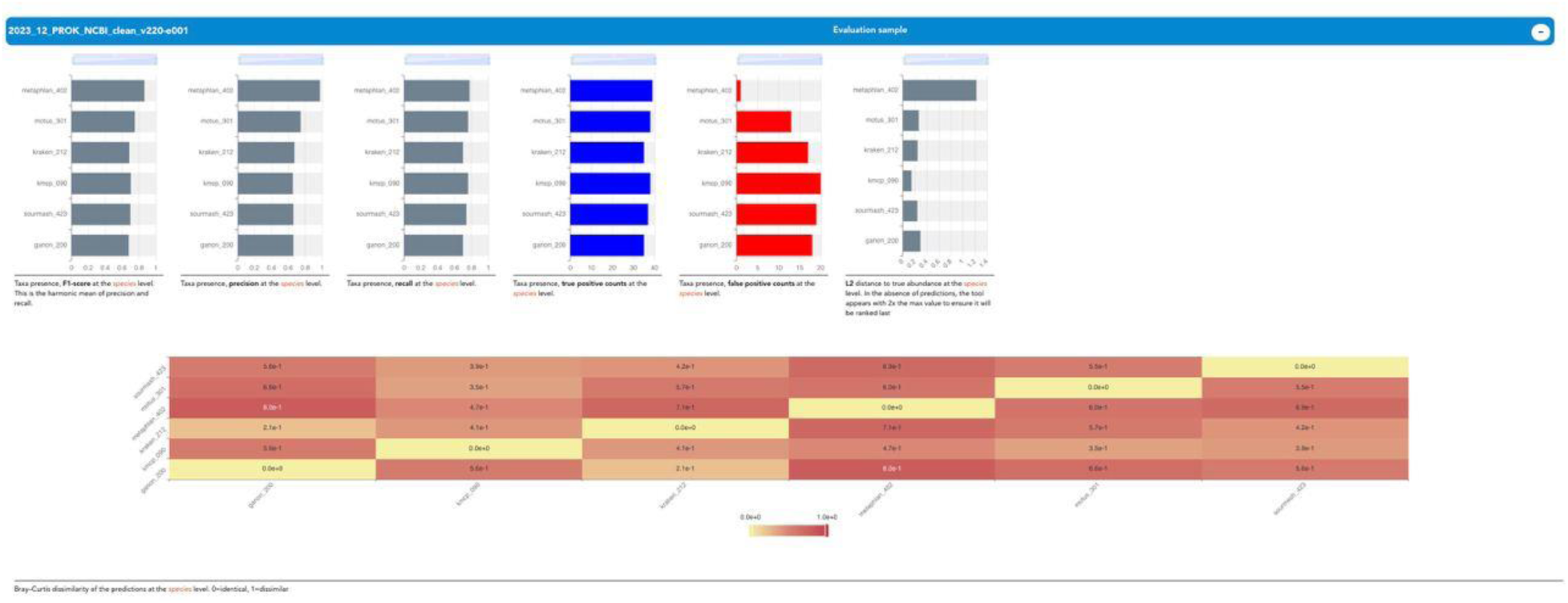
Detail page of a LEMMIv2 instance, showing different widgets reporting values for a specific evaluation sample, including a heatmap on how dissimilar the predictions of each tool are for that sample.

**Supplementary Figure 3.**
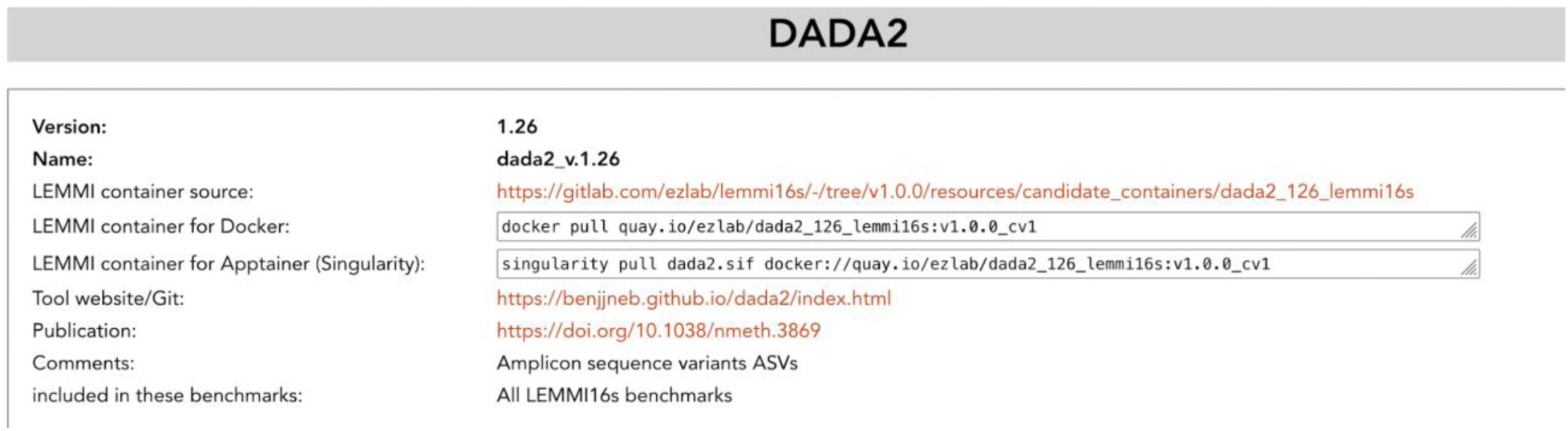
Extract from the LEMMI16S catalogue. Each tool has a section where different versions can be presented, with a link to the source of the container, the built container available in a repository, and links to a website and publication for the method where applicable.

**Supplementary Figure 4.**
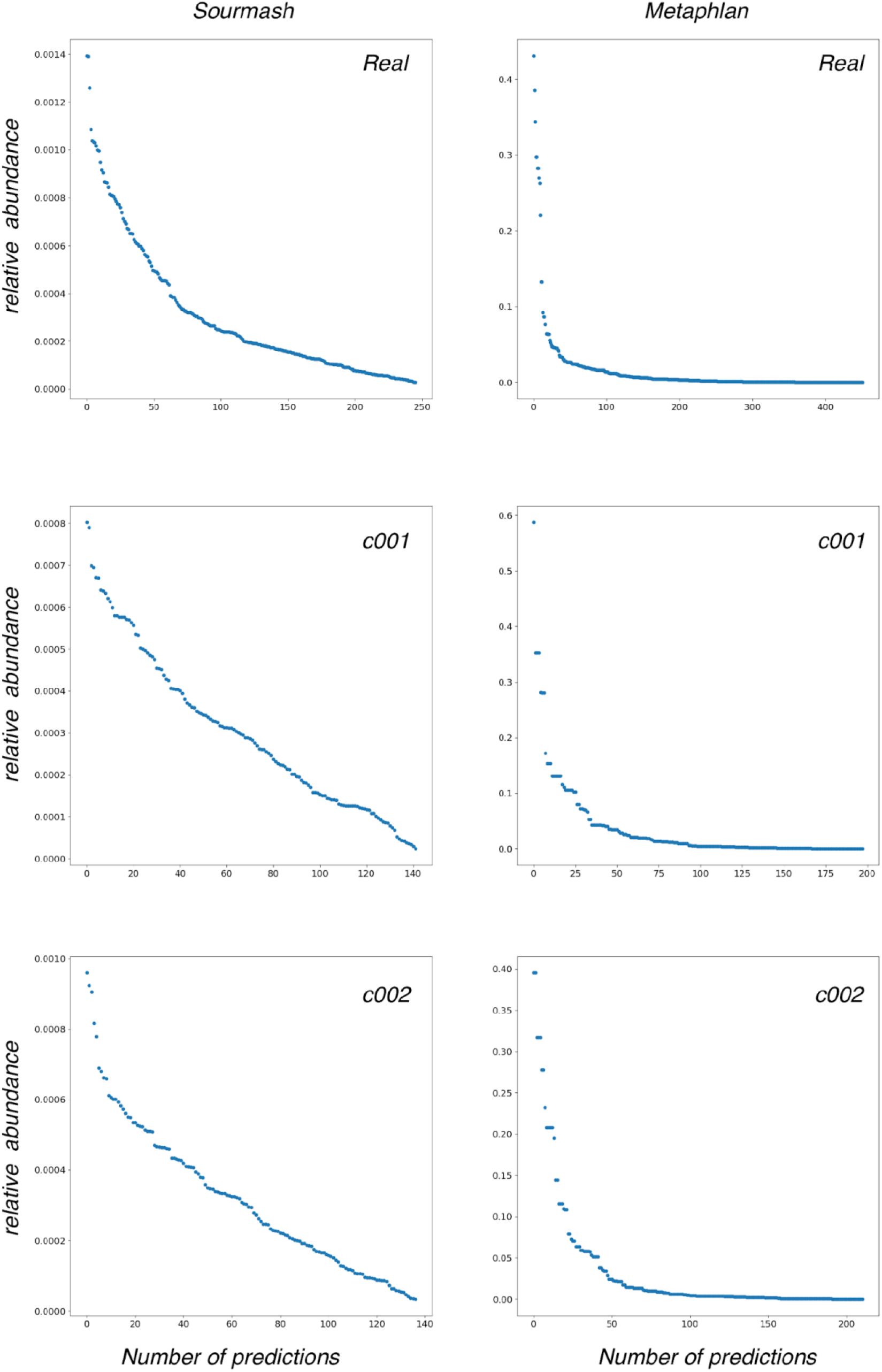
Relative abundances of taxa predicted by Sourmash and Metaphlan on LEMMIv2 samples 2023_05_PROK_NCBI-c001, c002 and the corresponding real sample. Includes predictions at all taxonomic ranks.

**Supplementary Figure 5.**
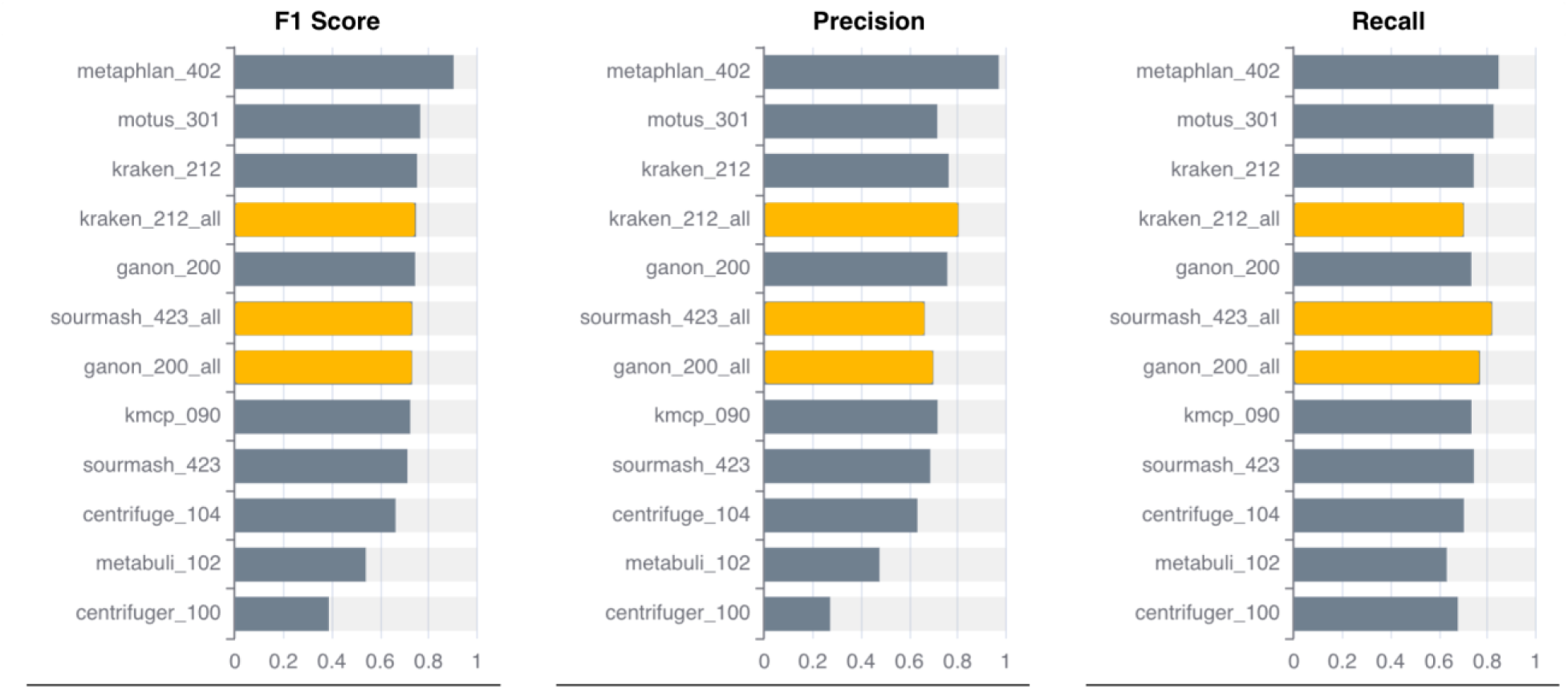
Performance of methods for the identification of organisms at species level for the instance 2023_12_PROK_NCBI_v220. Methods marked with _all used up to five genomes for each species as a reference. The others used only one genome as reference.

**Supplementary Figure 6.**
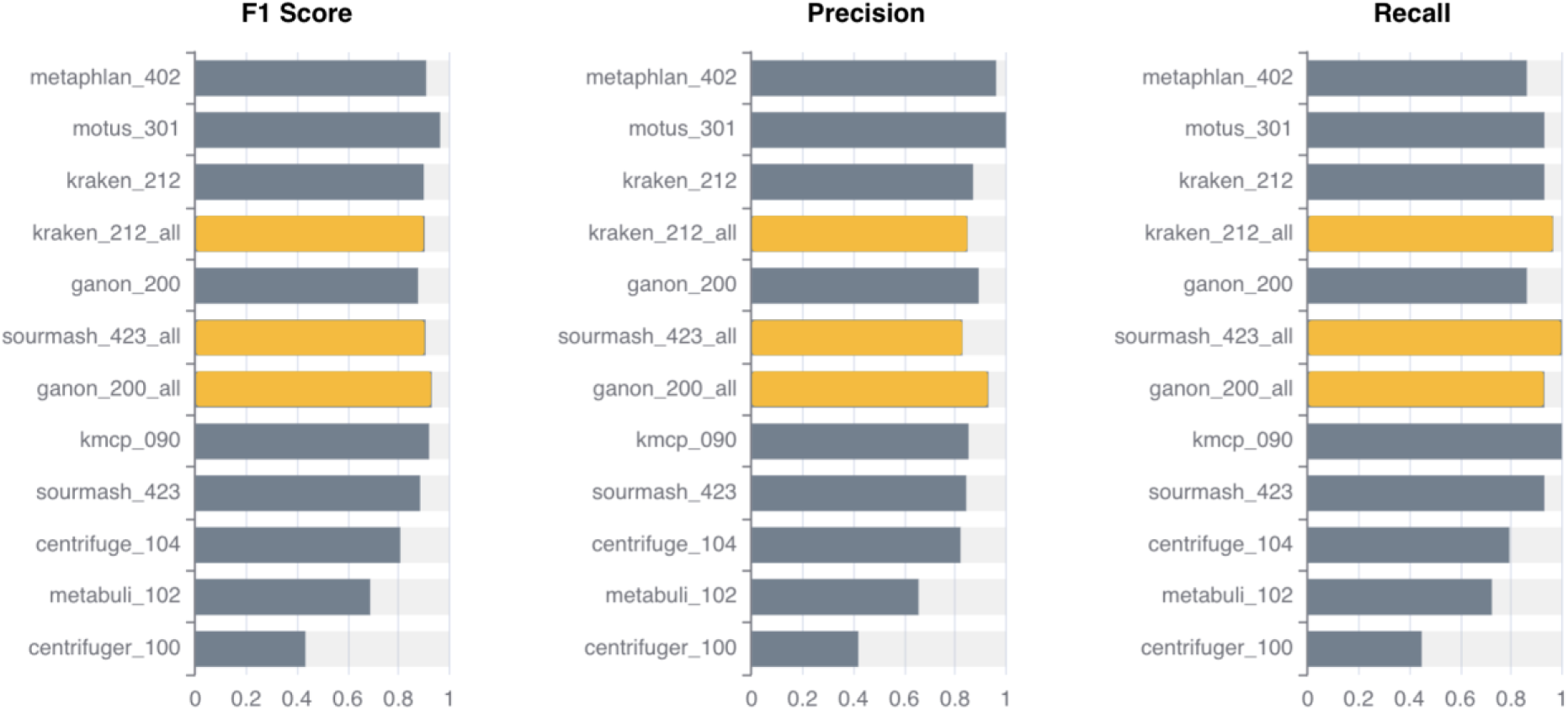
Performance of methods for the identification of organisms at genus level for the instance 2023_12_PROK_NCBI_v220. Methods marked with _all used up to five genomes for each species as a reference. The others used only one genome as reference.

**Supplementary Figure 7.**
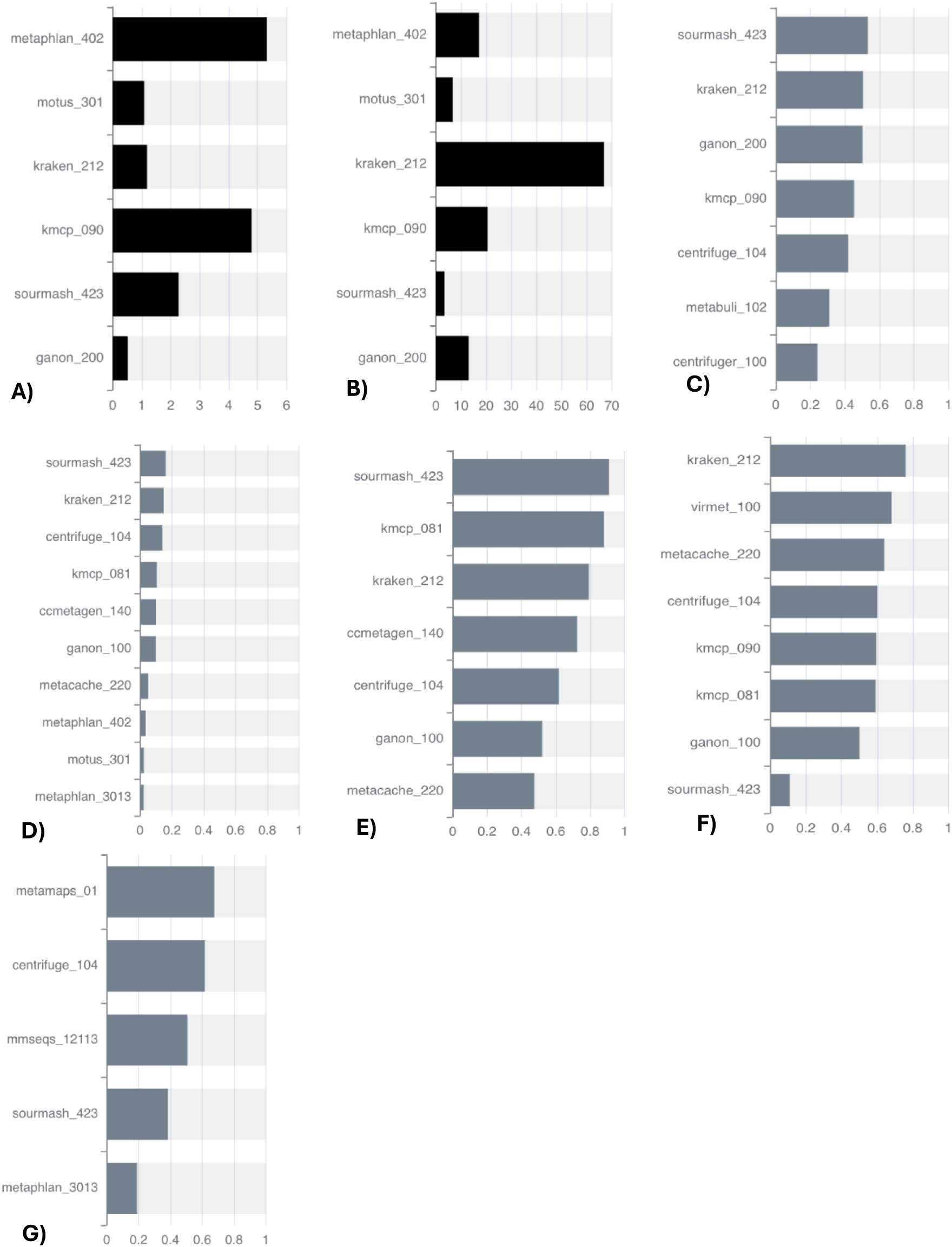
A) Average runtime (minutes) to perform the analysis of a sample from the 2023_12_PROK_NCBI_clean_v220 instance. B) Average peak memory (GB) to perform the analysis of a sample from the 2023_12_PROK_NCBI_clean_v220 instance. C) F1 score of methods for the identification of organisms at species level for the instance 2023_11_PROK_GTDB_v220. D) F1 score of methods for the identification of organisms at species level for the instance 2022_03_PROK_NCBI_2. E) F1 score of methods for the identification of organisms at species level for the instance 2022_01_EUK_NCBI. F) F1 score of methods for the identification of organisms at species level for the instance 2021_9_VIR_NCBI. G) F1 score of methods for the identification of organisms at species level for the instance 2022_04_PROK_NCBI_LR2.

**Supplementary Figure 8.**
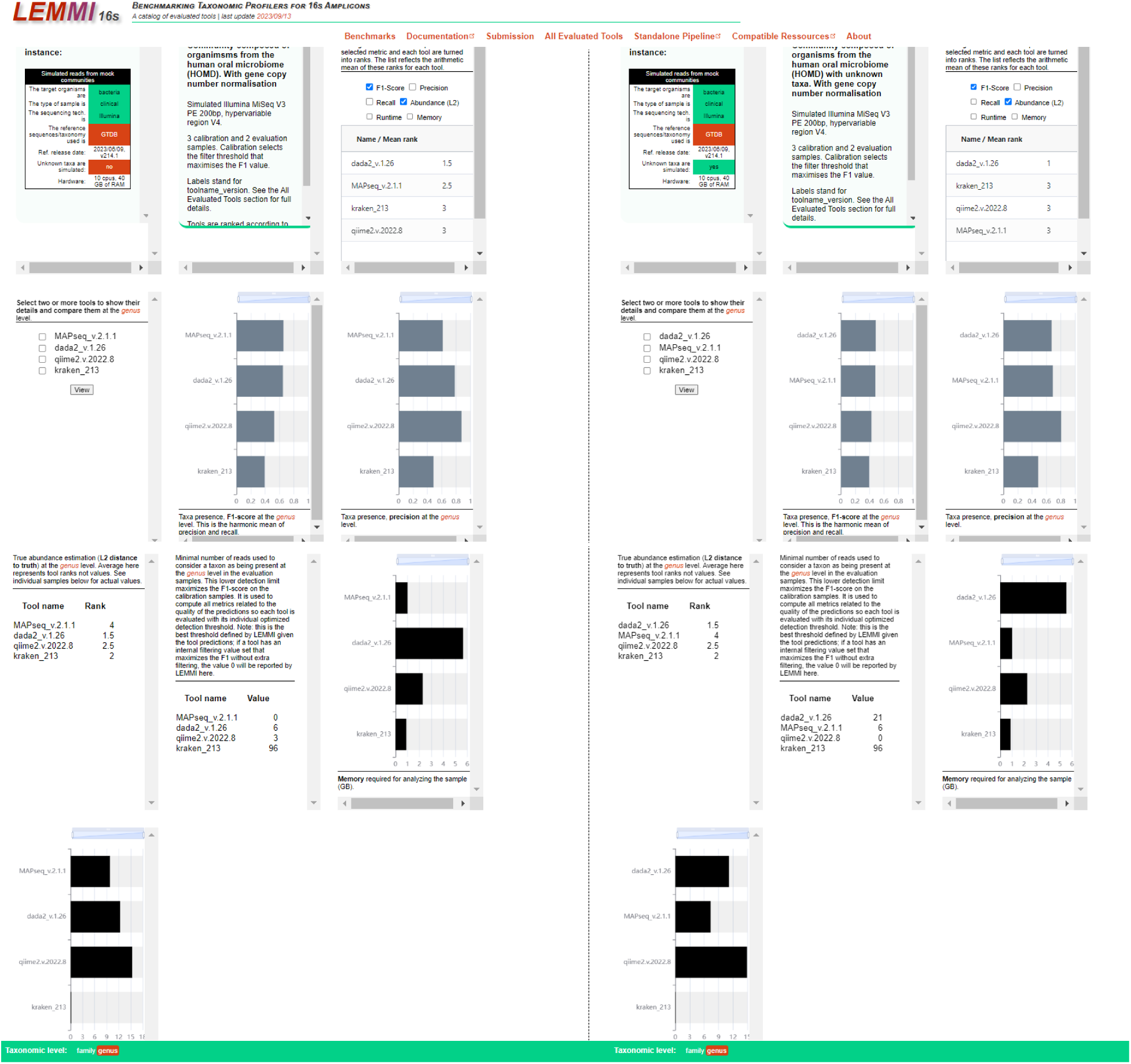
Results taken from LEMMI16S instance representing Human Oral Microbiome samples with unknown organisms.

**Supplementary Table 1.** Characteristics of simulated samples compared to the real samples on which they are based.

|  | Reads length | Number of paired-end reads | Number of 31-mers | %GC |
| --- | --- | --- | --- | --- |
| **LEMMIV2** |  |  |  |  |
| Plaque from a healthy patient | 151 | 21'153'159 | 2,059,607,399 | 45 |
| Simulation 2023_05_P ROK_NCBIC001 | 150 | 20'015'528 | 2,206,174,136 | 41 |
| Simulation 2023_05_P ROK_NCBIC002 | 150 | 20'055'700 | 2,224,328,630 | 41 |
| **LEMMI16S** |  |  |  |  |
| Human pathogens samples | 300 | 654,196 | 354,293,576 | 53.7 |
| Simulation Human pathogens samples_gc n-c001 | 250 | 867,086 | 380,472,649 | 52.4 |
| Simulation Human pathogens samples_gc n-c002 | 250 | 892,299 | 392,439,207 | 53.6 |

**Supplementary Table 2.** Status of the tools in terms of the ability of the container to filter out off-target reads prior to final classification. In some cases, the tool does not classify at the read level, making such pre-filtering impossible, and in other cases, the process was considered too heavy in terms of runtime and memory.

| Tool name | Filter host | Filter contaminants |
| --- | --- | --- |
| ccmetagen_140 | Yes | No |
| centrifuge_104 | Yes | Yes |
| centrifuge_100 | Yes | Yes |
| deepmicrobes_g | No | No |
| ganon_200, _160, _100 | Yes | Yes |
| kmcp_090, _081 | Yes | Yes |
| kraken_212 | Yes | Yes |
| metabuli_101 | Yes | Yes |
| metacache_220 | Yes | Yes |
| metamaps_01 | Yes | Yes |
| metaphlan_402, _3013 | No | No |
| mmseqs_12113 | Yes | Yes |
| motus_301 | No | No |
| sourmash_423 | Yes | Yes |
| virmet_100 | Yes | Yes |

**Supplementary Table 3.** Estimation of true abundance (L2 distance to truth) at species level. Average ranks in minimising error across all evaluation samples from instance 2023_12_PROK_NCBI_v220.

| Tool name | Value |
| --- | --- |
| metaphlan_402 | 12 |
| motus_301 | 3.25 |
| kraken_212 | 4.75 |
| kraken_212_all | 2.75 |
| ganon_200 | 6.5 |
| sourmash_423_all | 10 |
| ganon_200_all | 5.25 |
| kmcp_090 | 1.15 |
| sourmash_423 | 11 |
| centrifuge_104 | 4.5 |
| metabuli_102 | 7.75 |
| centrifuger_100 | 9 |

## Reference

Agustinho, D. P., Fu, Y., Menon, V. K., Metcalf, G. A., Treangen, T. J., & Sedlazeck, F. J. (2024). Unveiling microbial diversity: harnessing long-read sequencing technology. Nature methods, 1–13.

Allard, G., Ryan, F. J., Jeffery, I. B., & Claesson, M. J. (2015). SPINGO: a rapid species-classifier for microbial amplicon sequences. BMC Bioinformatics, 16, 1–8.

Beghini, F., McIver, L. J., Blanco-Míguez, A., Dubois, L., Asnicar, F., Maharjan, S., Mailyan, A., Manghi, P., Scholz, M., & Thomas, A. M. (2021). Integrating taxonomic, functional, and strain-level profiling of diverse microbial communities with bioBakery 3. elife, 10, e65088.

Blanco-Míguez, A., Beghini, F., Cumbo, F., McIver, L. J., Thompson, K. N., Zolfo, M., Manghi, P., Dubois, L., Huang, K. D., & Thomas, A. M. (2023). Extending and improving metagenomic taxonomic profiling with uncharacterized species using MetaPhlAn 4. Nature biotechnology, 41(11), 1633–1644.

Bolyen, E., Rideout, J. R., Dillon, M. R., Bokulich, N. A., Abnet, C. C., Al-Ghalith, G. A., Alexander, H., Alm, E. J., Arumugam, M., & Asnicar, F. (2019). Reproducible, interactive, scalable and extensible microbiome data science using QIIME 2. Nature biotechnology, 37(8), 852–857.

Brown, C. T., & Irber, L. (2016). sourmash: a library for MinHash sketching of DNA. Journal of open source software, 1(5), 27.

Buetas, E., Jordán-López, M., López-Roldán, A., D’Auria, G., Martínez-Priego, L., De Marco, G., Carda-Diéguez, M., & Mira, A. (2024). Full-length 16S rRNA gene sequencing by PacBio improves taxonomic resolution in human microbiome samples. BMC genomics, 25(1), 310.

Callahan, B. J., McMurdie, P. J., Rosen, M. J., Han, A. W., Johnson, A. J. A., & Holmes, S. P. (2016). DADA2: High-resolution sample inference from Illumina amplicon data. Nature methods, 13(7), 581–583.

Chen, T., Yu, W.-H., Izard, J., Baranova, O. V., Lakshmanan, A., & Dewhirst, F. E. (2010). The Human Oral Microbiome Database: a web accessible resource for investigating oral microbe taxonomic and genomic information. Database, 2010.

Cole, J. R., Wang, Q., Cardenas, E., Fish, J., Chai, B., Farris, R. J., Kulam-Syed-Mohideen, A., McGarrell, D. M., Marsh, T., & Garrity, G. M. (2009). The Ribosomal Database Project: improved alignments and new tools for rRNA analysis. Nucleic acids research, 37(suppl_1), D141–D145.

Dashper, S., Mitchell, H., Lê Cao, K.-A., Carpenter, L., Gussy, M., Calache, H., Gladman, S., Bulach, D., Hoffmann, B., & Catmull, D. (2019). Temporal development of the oral microbiome and prediction of early childhood caries. Scientific Reports, 9(1), 19732.

DeSantis, T. Z., Hugenholtz, P., Larsen, N., Rojas, M., Brodie, E. L., Keller, K., Huber, T., Dalevi, D., Hu, P., & Andersen, G. L. (2006). Greengenes, a chimera-checked 16S rRNA gene database and workbench compatible with ARB. Applied and environmental microbiology, 72(7), 5069–5072.

Dilthey, A. T., Jain, C., Koren, S., & Phillippy, A. M. (2019). Strain-level metagenomic assignment and compositional estimation for long reads with MetaMaps. Nature communications, 10(1), 3066.

Edgar, R. C. (2018). Accuracy of taxonomy prediction for 16S rRNA and fungal ITS sequences. PeerJ, 6, e4652.

Furstenau, T. N., Schneider, T., Shaffer, I., Vazquez, A. J., Sahl, J., & Fofanov, V. (2022). MTSv: rapid alignment-based taxonomic classification and high-confidence metagenomic analysis. PeerJ, 10, e14292.

Gihawi, A., Ge, Y., Lu, J., Puiu, D., Xu, A., Cooper, C. S., Brewer, D. S., Pertea, M., & Salzberg, S. L. (2023). Major data analysis errors invalidate cancer microbiome findings. MBio, 14(5), e01607–01623.

Glöckner, F. O., Yilmaz, P., Quast, C., Gerken, J., Beccati, A., Ciuprina, A., Bruns, G., Yarza, P., Peplies, J., & Westram, R. (2017). 25 years of serving the community with ribosomal RNA gene reference databases and tools. Journal of biotechnology, 261, 169–176.

Huang, W., Li, L., Myers, J. R., & Marth, G. T. (2012). ART: a next-generation sequencing read simulator. Bioinformatics, 28(4), 593–594.

Huerta-Cepas, J., Serra, F., & Bork, P. (2016). ETE 3: reconstruction, analysis, and visualization of phylogenomic data. Molecular biology and evolution, 33(6), 1635–1638.

Johnson, J. S., Spakowicz, D. J., Hong, B.-Y., Petersen, L. M., Demkowicz, P., Chen, L., Leopold, S. R., Hanson, B. M., Agresta, H. O., & Gerstein, M. (2019). Evaluation of 16S rRNA gene sequencing for species and strain-level microbiome analysis. Nature communications, 10(1), 5029.

Kim, D., Song, L., Breitwieser, F. P., & Salzberg, S. L. (2016). Centrifuge: rapid and sensitive classification of metagenomic sequences. Genome research, 26(12), 1721–1729.

Kim, J., & Steinegger, M. (2024). Metabuli: sensitive and specific metagenomic classification via joint analysis of amino acid and DNA. Nature methods, 1–3.

Köster, J., & Rahmann, S. (2012). Snakemake—a scalable bioinformatics workflow engine. Bioinformatics, 28(19), 2520–2522.

Kurtzer, G. M., Sochat, V., & Bauer, M. W. (2017). Singularity: Scientific containers for mobility of compute. PLoS One, 12(5), e0177459.

Liang, Q., Bible, P. W., Liu, Y., Zou, B., & Wei, L. (2020). DeepMicrobes: taxonomic classification for metagenomics with deep learning. NAR Genomics and Bioinformatics, 2(1), lqaa009.

López-Aladid, R., Fernández-Barat, L., Alcaraz-Serrano, V., Bueno-Freire, L., Vázquez, N., Pastor-Ibáñez, R., Palomeque, A., Oscanoa, P., & Torres, A. (2023). Determining the most accurate 16S rRNA hypervariable region for taxonomic identification from respiratory samples. Scientific Reports, 13(1), 3974.

Mangul, S., Martin, L. S., Hill, B. L., Lam, A. K.-M., Distler, M. G., Zelikovsky, A., Eskin, E., & Flint, J. (2019). Systematic benchmarking of omics computational tools. Nature communications, 10(1), 1393.

Marcelino, V. R., Clausen, P. T., Buchmann, J. P., Wille, M., Iredell, J. R., Meyer, W., Lund, O., Sorrell, T. C., & Holmes, E. C. (2020). CCMetagen: comprehensive and accurate identification of eukaryotes and prokaryotes in metagenomic data. Genome biology, 21, 1–15.

Matias Rodrigues, J. F., Schmidt, T. S., Tackmann, J., & von Mering, C. (2017). MAPseq: highly efficient k-mer search with confidence estimates, for rRNA sequence analysis. Bioinformatics, 33(23), 3808–3810.

McIntyre, A. B., Ounit, R., Afshinnekoo, E., Prill, R. J., Hénaff, E., Alexander, N., Minot, S. S., Danko, D., Foox, J., & Ahsanuddin, S. (2017). Comprehensive benchmarking and ensemble approaches for metagenomic classifiers. Genome biology, 18, 1–19.

Müller, A., Hundt, C., Hildebrandt, A., Hankeln, T., & Schmidt, B. (2017). MetaCache: context-aware classification of metagenomic reads using minhashing. Bioinformatics, 33(23), 3740–3748.

Nasko, D. J., Koren, S., Phillippy, A. M., & Treangen, T. J. (2018). RefSeq database growth influences the accuracy of k-mer-based lowest common ancestor species identification. Genome biology, 19, 1–10.

O’Sullivan, D. M., Doyle, R. M., Temisak, S., Redshaw, N., Whale, A. S., Logan, G., Huang, J., Fischer, N., Amos, G. C., & Preston, M. D. (2021). An inter-laboratory study to investigate the impact of the bioinformatics component on microbiome analysis using mock communities. Scientific Reports, 11(1), 10590.

Parks, D. H., Chuvochina, M., Waite, D. W., Rinke, C., Skarshewski, A., Chaumeil, P.-A., & Hugenholtz, P. (2018). A standardized bacterial taxonomy based on genome phylogeny substantially revises the tree of life. Nature biotechnology, 36(10), 996–1004.

Piro, V. C., Dadi, T. H., Seiler, E., Reinert, K., & Renard, B. Y. (2020). ganon: precise metagenomics classification against large and up-to-date sets of reference sequences. Bioinformatics, 36(Supplement_1), i12–i20.

Piro, V. C., & Reinert, K. (2023). ganon2: up-to-date and scalable metagenomics analysis. bioRxiv, 2023.2012. 2007.570547.

Pruesse, E., Quast, C., Knittel, K., Fuchs, B. M., Ludwig, W., Peplies, J., & Glöckner, F. O. (2007). SILVA: a comprehensive online resource for quality checked and aligned ribosomal RNA sequence data compatible with ARB. Nucleic acids research, 35(21), 7188–7196.

Robeson, M. S., O’Rourke, D. R., Kaehler, B. D., Ziemski, M., Dillon, M. R., Foster, J. T., & Bokulich, N. A. (2021). RESCRIPt: Reproducible sequence taxonomy reference database management. PLoS computational biology, 17(11), e1009581.

Ruscheweyh, H.-J., Milanese, A., Paoli, L., Karcher, N., Clayssen, Q., Keller, M. I., Wirbel, J., Bork, P., Mende, D. R., & Zeller, G. (2022). Cultivation-independent genomes greatly expand taxonomic-profiling capabilities of mOTUs across various environments. Microbiome, 10(1), 212.

Saini, J. S., Hassler, C., Cable, R., Fourquez, M., Danza, F., Roman, S., Tonolla, M., Storelli, N., Jacquet, S., & Zdobnov, E. M. (2022). Bacterial, phytoplankton, and viral distributions and their biogeochemical contexts in meromictic Lake Cadagno offer insights into the Proterozoic Ocean microbial loop. MBio, 13(4), e00052–00022.

Schloss, P. D., & Handelsman, J. (2005). Introducing DOTUR, a computer program for defining operational taxonomic units and estimating species richness. Appl Environ Microbiol, 71(3), 1501–1506. 10.1128/AEM.71.3.1501-1506.2005

Schloss, P. D., Westcott, S. L., Ryabin, T., Hall, J. R., Hartmann, M., Hollister, E. B., Lesniewski, R. A., Oakley, B. B., Parks, D. H., & Robinson, C. J. (2009). Introducing mothur: open-source, platform-independent, community-supported software for describing and comparing microbial communities. Applied and environmental microbiology, 75(23), 7537–7541.

Schoch, C. L., Ciufo, S., Domrachev, M., Hotton, C. L., Kannan, S., Khovanskaya, R., Leipe, D., Mcveigh, R., O’Neill, K., & Robbertse, B. (2020). NCBI Taxonomy: a comprehensive update on curation, resources and tools. Database, 2020, baaa062.

Seppey, M., Manni, M., & Zdobnov, E. M. (2020). LEMMI: a continuous benchmarking platform for metagenomics classifiers. Genome research, 30(8), 1208–1216.

Shen, W., Xiang, H., Huang, T., Tang, H., Peng, M., Cai, D., Hu, P., & Ren, H. (2023). KMCP: accurate metagenomic profiling of both prokaryotic and viral populations by pseudo-mapping. Bioinformatics, 39(1), btac845.

Song, L., & Langmead, B. (2024). Centrifuger: lossless compression of microbial genomes for efficient and accurate metagenomic sequence classification. Genome biology, 25(1), 106.

Steinegger, M., & Söding, J. (2017). MMseqs2 enables sensitive protein sequence searching for the analysis of massive data sets. Nature biotechnology, 35(11), 1026–1028.

Weging, S., Gogol-Döring, A., & Grosse, I. (2021). Taxonomic analysis of metagenomic data with kASA. Nucleic acids research, 49(12), e68–e68.

Wood, D. E., Lu, J., & Langmead, B. (2019). Improved metagenomic analysis with Kraken 2. Genome biology, 20, 1–13.

Yang, C., Chu, J., Warren, R. L., & Birol, I. (2017). NanoSim: nanopore sequence read simulator based on statistical characterization. Gigascience, 6(4), gix010.

Zhou, N., Zhou, H., & Hoppe, D. (2022). Containerization for high performance computing systems: Survey and prospects. IEEE Transactions on Software Engineering, 49(4), 2722–2740.

